# A general mathematical framework for modelling subnetworks of the nuclear auxin pathway

**DOI:** 10.64898/2026.08.06.742982

**Authors:** Joseph G. Shuttleworth, Emily Chan, Thomas Welch, Rahul Bhosale, Anthony Bishopp, Etienne Farcot

## Abstract

Auxins are a family of plant hormones involved in various processes across plant tissues and species. The Nuclear Auxin Pathway (NAP) consists of interacting transcription factors (ARFs) and repressors (Aux/IAAs), which govern an individual cell’s response to changes in auxin concentration. These components are present in all land plants, and many species possess multiple copies of each signalling component. We present a general framework for ODE-based models of NAP submodules with the flexibility to model the promotion and repression of target genes by any combination of transcriptional regulators. We analyse published data and show that auxin treatment in *Arabidopsis thaliana* roots triggers a range of characteristically distinct temporal response profiles—for both target genes and the signalling components themselves. Using our modelling framework, we recapitulate aspects of this behaviour by presenting examples of real and theoretical NAP subnetworks, and by analysing the effect that these network dynamics have on auxin-mediated transcriptional responses. This work demonstrates the utility of our modelling framework as a general-purpose tool for understanding the function of certain protein-protein and protein-DNA interactions through their effects on the NAP. This exploration of the rich dynamics of more complex signalling pathways promises to advance our understanding of the NAP.

## 1 Introduction

Auxins comprise a family of plant hormones that are implicated in many processes across plant species. An individual cell’s response to changes in auxin concentration is governed by a signal transduction pathway referred to as the *Nuclear Auxin Pathway* (NAP). The main components of this regulatory network are known to be auxin/indole acetic acid repressor proteins (Aux/IAA), and transcription factors known as Auxin Response Factors (ARFs), and proteins from the TIR1/AFB (TRANSPORT INHIBITOR RESPONSE1/AUXIN SIGNALING F-BOX) family. Aux/IAAs and ARFs contain similar domains (domains III/IV) that allow protein-protein binding [47]. Through the binding of these domains, Aux/IAAs and ARFs combine to form multimers, complexes containing a combination of ARFs and Aux/IAAs. Such multimers may contain ARFs, Aux/IAAs or a mixture of proteins from both families. Certain multimers containing ARFs bind to the cell’s DNA at an *AuxRE* (auxin-response element), up/down regulating the expression of specific genes, such as those in the small-auxin upregulated RNA (SAUR) family [17]. Specific ARFs are known to promote the transcription of auxin-related genes, whilst others are known to repress transcription. Aux/IAAs do not bind directly to the DNA themselves but can bind via dimerisation with ARFs and repress transcription by recruiting TOPLESS (TPL) proteins [49].

Auxin interacts with this system by mediating the binding of TIR1/AFB and Aux/IAA proteins, which results in the ubiquitination and degradation of Aux/IAA via the SCF-type ubiquitin ligase complex [42]. This removal of Aux/IAA decreases competition, increasing the probability that an ARF-ARF dimer is bound to the *AuxRE*. The exact effect of changes in auxin concentration is therefore dependent on the specific *signalling components* (ARFs, Aux/IAAs and TIR1/AFB proteins) present in the nucleus, their concentrations, and their propensity for binding to each other and relevant promoter regions in the DNA [27]. Figure 1 depicts the role of each component in the NAP, and how changes in auxin concentration drive transcriptional regulation of target genes.

**Figure 1.**
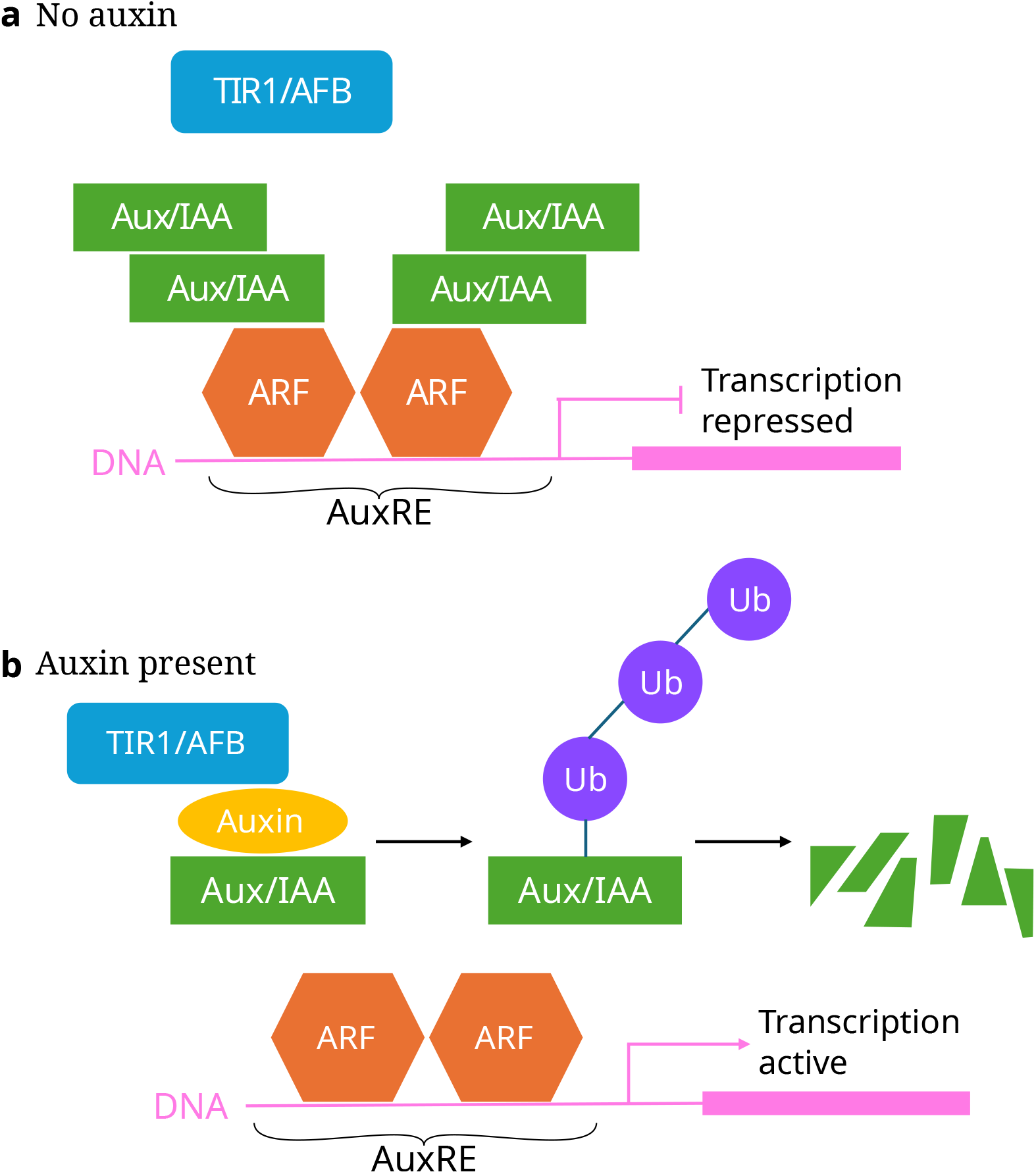
The canonical mechanism by which the NAP causes a transcriptional response in target genes. **a** When there is little or no auxin, Aux/IAAs are free to bind to ARFs (which themselves bind to the DNA) and repress transcription of the target gene. **b**: Auxin causes the ubiquitination and subsequent decay of Aux/IAAs, freeing promoter ARFs to up-regulate the target gene.

In the simplest instance, as the auxin concentration increases, we may expect (at steady state) the concentration of Aux/IAAs and mRNA for particular target genes, such as the family of small auxin-upregulated RNA (SAUR) genes, to increase [36]. However, the complexity of the NAP means that there is a rich set of possible dynamics (including multistability and oscillations, for example) dictating the response of a given cell to a specific auxin stimulus. The NAP can be seen as a large, general-purpose network containing every signalling component encoded for in the genome, and, simultaneously, as a family of *subnetworks*, or modules, where only certain cell-specific combinations of signalling components are highly expressed [44, 48, 32].

Whilst these signalling components are present in all multicellular plants, each protein family (TIR1/AFB, Aux/IAA, ARF) is larger in seed plants. This additional complexity seems to coincide with a more diverse range of cell-specific auxin responses that in early diverging land plants [24, 23, 32]. The NAP of the model plant species, *Arabidopsis thaliana*, consists of multiple distinct proteins in each of these families: 6 TIR1/AFB proteins, 29 Aux/IAAs, and 23 ARFs of the respective components [12]. In comparison, the NAP of *Marchantia polymorpha* (common liverwart), consists of 1 TIR1/AFB protein, 1 Aux/IAA and 3 distinct ARF proteins. With fewer components, the NAP system of *Marchantia polymorpha* facilitates key developmental processes [13], but does not exhibit the same degree of cell-specificity as *Arabidopsis thaliana* [3].

In general, mathematical models aid our understanding of molecular pathways, such as the NAP, by showing how individual regulatory mechanisms interact to modulate cellular development and other biological processes, thereby elucidating the role of regulatory elements [20]. These trans-regulatory elements themselves are also regulated by auxin, meaning that NAP subnetworks contain multiple positive and negative feedback loops, which may give rise to highly nonlinear dynamics [27, 16]. In such systems, even when the properties of individual components are well studied, their function within the context of the network as a whole may be unintuitive. Building accurate mathematical models of the NAP is therefore important for achieving a functional understanding of auxin signalling. Here, we introduce a general mathematical framework that can be applied to any NAP subnetwork in any species.

In Section 2, we analyse transcriptomic data previously collected from *Arabidopsis thaliana*, characterising the response of various genes to an exogenously applied auxin treatment. The diversity of temporal response profiles found across the three ARF classes further motivates the introduction of a general NAP modelling framework, while providing an experimental test-case for its validation. Using this framework, we demonstrate that with only a few interacting signalling components, a diverse range of auxin-mediated transcriptional responses can be found.

Numerous mathematical models of specific instances of the NAP have been proposed previously [29, 45, 2, 5, 11]. However, these models typically describe only a single ARF and a single Aux/IAA. Moving beyond single-ARF, single-Aux/IAA models, Bridge et. al [5] propose a model with two ARFs and two Aux/IAAs, though it is assumed that ARF concentrations are constant, and, as such, this model is unable to recapitulate the transcriptional regulation of the ARFs themselves. In Section 3 we present a flexible modelling framework that can be used to model instances of the NAP with multiple ARFs and Aux/IAAs. Given that transcriptional responses to auxin are diverse across plant species and cell types, we present our model as a flexible, general-purpose tool that can be applied to any NAP with various combinations of signalling components (namely, ARFs and Aux/IAAs). We also provide Julia scripts for all the simulations and analyses in this paper. With simple modifications, these codes may be used to perform similar analyses for any model within our general framework.

Using this framework, we present three specific examples of network motifs. The first, in Section 44.1, concerns monopteros/ARF5, a key ARF which is commonly expressed across cell types in *Arabidopsis thaliana*, and bodenlos/IAA12 (BDL), which is known to interact with monopteros (MP). Here, we show that even in a simple model with a single ARF and Aux/IAA, non-trivial dynamics and bistability are possible. Then, in Section 44.2, we add a repressor ARF to this example, and show a tipping-point mechanism, where excessive repressive activity switches off transcriptional activity, reducing the expression of other auxin-responsive genes (including auxin-signalling components).

In Section 44.3, we explore a hypothetical but plausible model consisting of a single Aux/IAA and two interacting activator ARFs, with the first regulating the expression of the second. In *Arabidopsis thaliana*, ARF7 and ARF19 are a well-studied pair of activator ARFs that provide an example of such a module [16]. ARF19 is promoted by ARF7 and acts to amplify the promotion of specific target genes [16], reminiscent of the well-studied feedforward motif [26]. To capture transcriptional responses at different timepoints, we include additional self-promotional feedback loop for the expression of the downstream ARF. This example network serves to demonstrate plausible functional outputs arising from interacting promoter ARFs. In particular, we show how model output is sensitive to specific parameters relating to protein-protein interactions, and how these parameters can induce delays in target-gene transcription and allow frequency-sensitive responses to oscillatory signals.

## 2 Auxin-responsive genes show transcriptional responses across different timescales

Experimental studies have shown that the auxin response is replete with feed-forward and feed-back loops that provide subnetworks with specificity in auxin response [27, 44, 48, 32]. To anchor our modelling within a realistic framework of auxin response dynamics, we re-analysed previous data to examine the timing of response dynamics to an auxin treatment in a single tissue type.

It has been observed that auxin-responsive genes are up-/down-regulated at different times in response to different auxin treatments in different cell types [34]. However, analyses relying on few observation times may misrepresent temporal expression profiles, leading to a mischaracterisation of transcriptional response.

Here, we explore timecourse microarray data for exogenous application of indole-3-acetic acid (IAA), a naturally occurring plant auxin, on *Arabidopsis thaliana* root meristems [50, 46]. We first summarise the genome-wide transcriptional responses found, before focusing on the transcription of certain trans-regulatory elements (namely ARFs and Aux/IAAs), given their key roles in regulating the NAP.

Microarray technology allows high-throughput gene expression profiling by quantifying transcripts present in a sample through hybridization of the corresponding fluorescently-labelled complementary DNA (cDNA) to chips containing thousands of DNA probes for known transcripts in the target genome [1].

Figure 2 highlights different timescales of transcriptional responses modulated by auxin, from the genes showing differential expression with at least four-fold increase or decrease in gene expression level, that is, where the log_2_-fold change is greater than 2 or less than −2. Strikingly, some genes are specifically highly up-regulated within 30 minutes of IAA treatment, and subsequently decrease in expression levels at later time points—such genes are likely primary targets of the auxin signalling pathway [34]. We also observe other genes that are only up-regulated at later time points of IAA treatment, and also genes that are down-regulated (either rapidly by the first 30-minute time point, or more slowly over two hours). These data show that within this tissue type that there is no stereotypical response, and rather clusters of genes respond with specific temporal profiles. We hypothesise that this could be due to different clusters of genes being regulated by different subnetworks.

**Figure 2.**
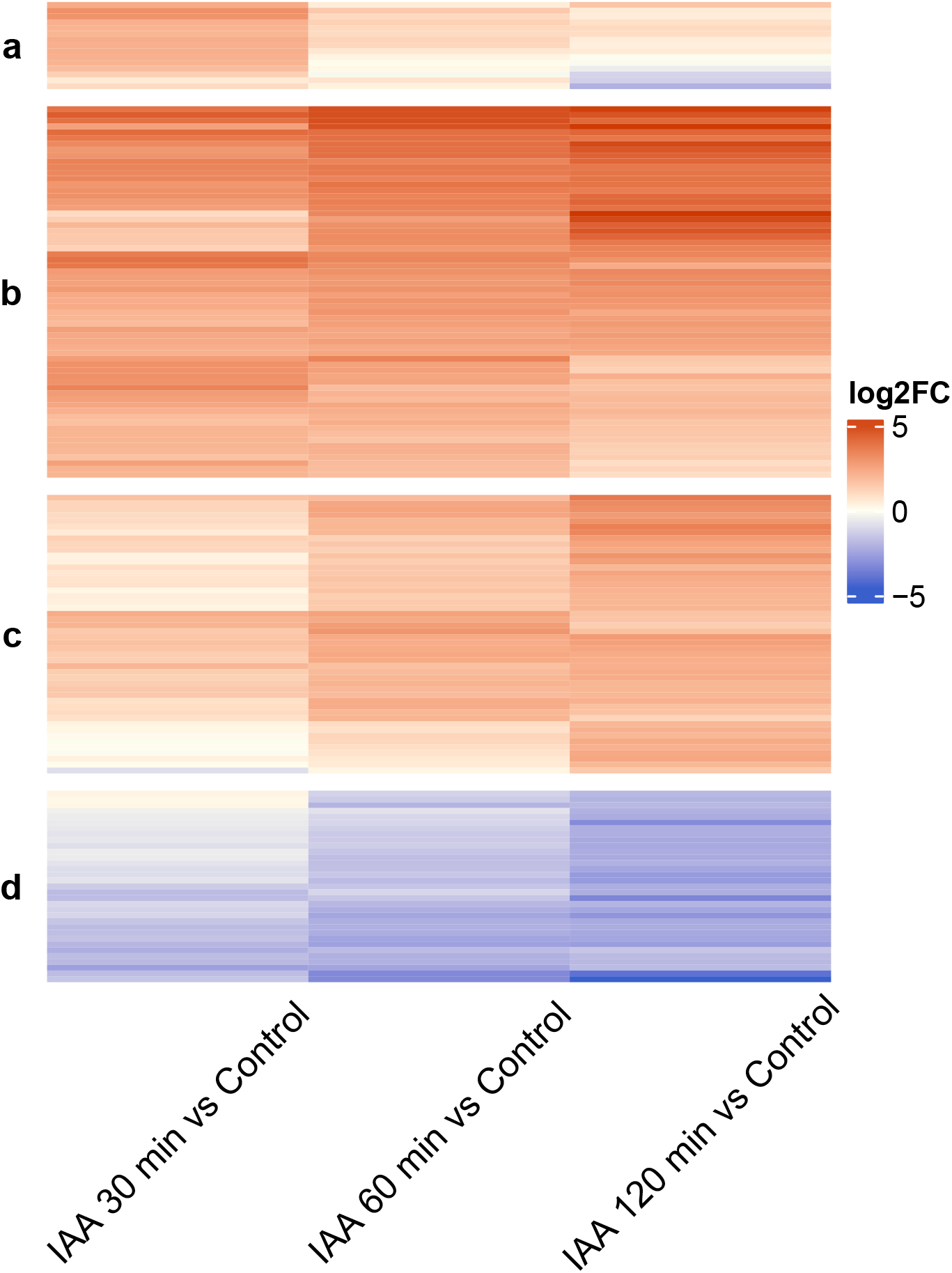
Heatmaps of temporal profiles of gene expression in response to IAA treatment in *Arabidopsis thaliana* root meristems [46]. Genes with a relative differential expression greater than 4 (that is, a log_2_-fold change greater than 2) are clustered according to their response profile using Euclidean distance and complete linkage. Each row shows the log_2_-fold change (**log2FC**) for a given gene at different time points of IAA treatment relative to the control (no IAA). A positive log_2_-fold change (red shades in colour bar) denotes upregulation of a given gene for a given duration of IAA treatment, whereas negative log_2_-fold change (blue shades in colour bar) denotes downregulation of a given gene. Genes clustered by log_2_-fold change across timepoints roughly fall into four groups showing up-/down-regulation on different timescales. Group **a**: Genes which are rapidly upregulated at 30 minutes, but for which gene expression falls at later time points. Group **b**: Genes which are consistently upregulated across all time points. Group **c**: Genes which are slowly upregulated— often showing little upregulation at 30 minutes, but which become more upregulated at later time points. Group **d**: downregulated genes: some of which are downregulated rather slowly, similar to the delayed upregulation exhibited by group **c**.

To see whether the diversity of temporal auxin responses could be correlated with changes in auxin signalling components, we examined how NAP components changed in this dataset.

Figure 2 shows the transcriptional response for a selection of genes (those meeting a *±*2 log_2_-fold change in expression level relative to the untreated control). Here, we see that different target genes have characteristically different responses, with some genes being upregulated and others being downregulated, both at various timepoints.

Figure 3(**b**) shows the log_2_-fold changes in expression levels of the Aux/IAAs present in the dataset. Here, we observe that many Aux/IAAs are up-regulated rapidly (within 30 minutes) in response to auxin, with only a small number seemingly being unaffected or only very slightly down-regulated. There is also diversity in these temporal expression patterns, with some Aux/IAAs, such as IAA29 and IAA30, increasing in expression throughout the experiment. However, we do not see the same variety of up– and down-regulation as with the ARF transcripts shown in Figure 3(**a**). Full computer codes for the production the post-processing of the microarray data and these calculations are found in the Supplementary Materials.

**Figure 3.**
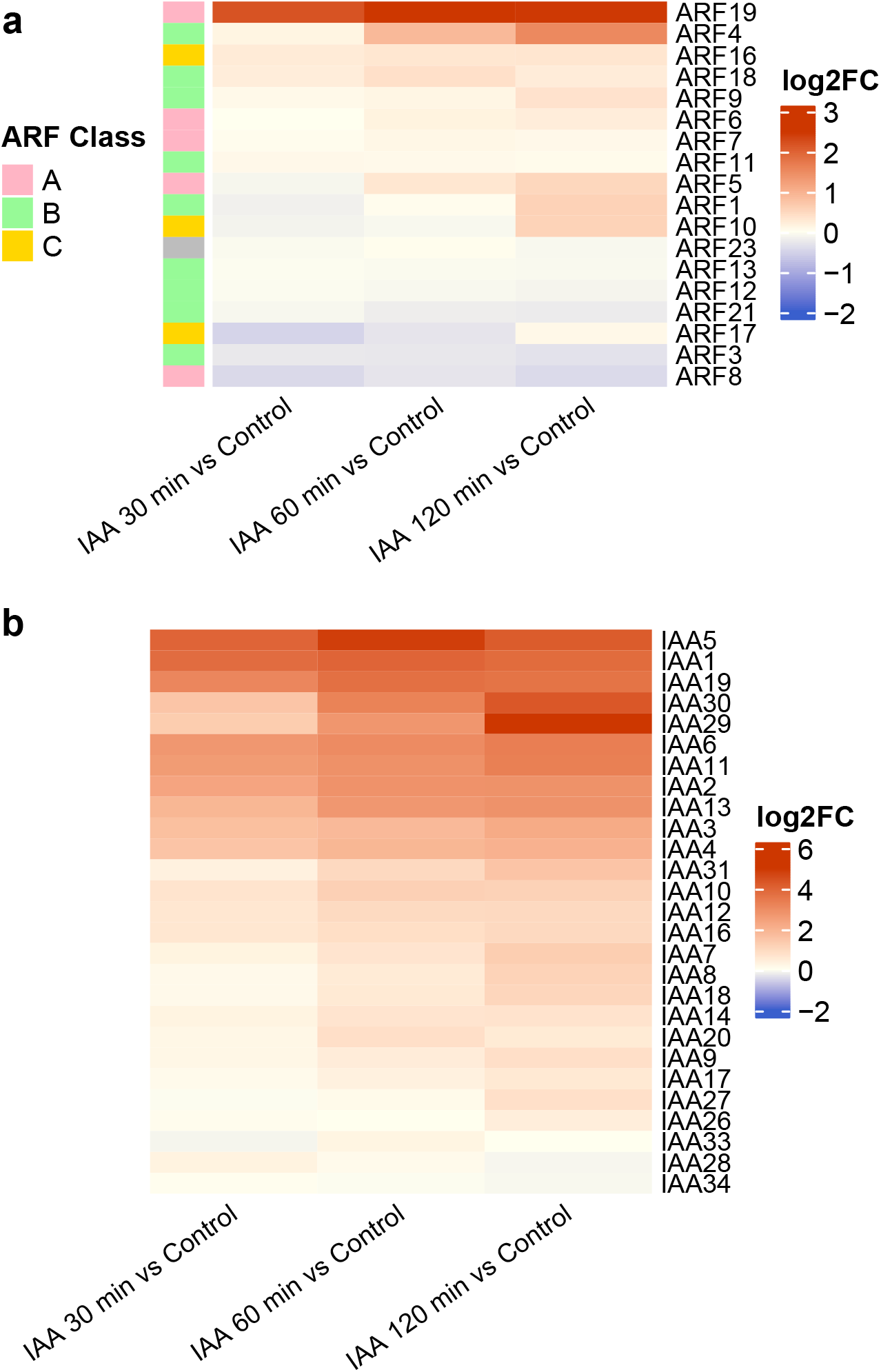
Heatmaps showing relative changes in expression level of *Arabidopsis thaliana* auxin-signalling components after auxin (IAA) treatment. (**a**): log_2_ fold-changes in expression of ARFs after IAA treatment compared to a control (no IAA). Genes are clustered according to their response profiles using Euclidean distance and complete linkage. The colours on the left of the heatmap show the class (that is, A, B, or C) of ARF that a given transcript belongs too (ARF23 is coloured grey as it is denoted a pseudogene [8]). Note how a range of different expression patterns can be seen within each ARF class. **b**: log_2_ fold-changes in Aux/IAA expression after IAA treatment compared to a control (no IAA). Whilst some Aux/IAAs (IA7 A28 and IAA34) are slightly down-regulated, the log_2_-fold change is much greater than −2 (a typical cutoff value).

These data examine one tissue type in a single plant species, at one developmental stage, and treated with a single auxin concentration. Nevertheless, we observe distinct response profiles amongst ARFs and Aux/IAAs. It is highly likely that we are only capturing a small proportion of the total feedbacks within the whole plant kingdom. Therefore, we envision that developing mathematical models incorporating multiple ARFs and Aux/IAAs, where each component can have distinct modes of regulation, will provide a platform to explore the complexity of the auxin response.

To better understand the emergent behaviour arising from such interactions, we propose a mathematical modelling framework that has the flexibility to include any of these possible regulatory mechanisms. Because of the diversity of transcriptional responses exhibited by ARFs described above, we focus on simple subnetworks with multiple ARF species which regulate each other’s expression. Through these examples, we demonstrate that a rich array of dynamics are possible in small subnetworks and individual motifs of the NAP.

## 3 Mathematical modelling of the auxin signalling pathway

### 3.1 Single-ARF, Single-Aux/IAA model

Many models of auxin signalling exist, and we build upon these by developing a flexible framework for incorporating multiple NAP components with distinct regulations. On the molecular scale, the NAP consists of many stochastic interactions between individual components in the nucleus. As in previously proposed models [29, 45, 2, 5, 11], we approximate the overall effect of such interactions using a *mass-action* approximation, where we consider only the amount/concentration of each protein/compound of interest. We model the NAP using a system of Ordinary Differential Equations (ODEs), each of which describes the concentration of a given species in a single cell (that is, the concentration of mRNA for a given gene or the concentration of a given protein, for example).

Here, we make a number of further simplifying assumptions. Firstly, for the sake of parsimony, we consider only monomers and dimers and not multimers of higher order. Whilst the formation of large multimers have been observed experimentally [33, 38], for simplicity, we assume that the overall dynamics of the system can be described by only considering protein complexes with up to two components.

Specific domains mediate protein-protein binding in ARFs (Domains III and IV) and Aux/IAAs (PB1). Through these domains, pairs of these components bind with each other forming *dimers*— ARF-ARF, ARF-Aux/IAA and Aux/IAA-Aux/IAA dimers may all be formed [44]. This process, dimerisation, has important implications. For example, Aux/IAAs in the vicinity of the promoter region recruit the transcriptional co-repressor, TPL, repressing transcription. The different ARF species have different preferences and affinities for binding with the various motifs found in each promoter region [28]. Moreover, once such a dimer is bound to the promoter, the combinations of two different activation domains (ADs) may cause a specific transcriptional response.

So that we can explore the possible effects of dimerisation on the NAP’s transcriptional response, we explicitly include the concentration of such dimers in our model as state variables. We denote the concentration of a dimer formed by a single ARF and Aux/IAA as *D_A,I_*, with the subscript *A* and *I* denoting the ARF and Aux/IAA, respectively. For simplicity, we assume that the order of these dimers are inconsequential such that *D_A,I_* = *D_I,A_*, for example. In this way, the only dimer concentrations we need to consider are *D_A,A_, D_A,I_*, and *D_I,I_*. We denote the ARF and Aux/IAA mRNA concentration by *R_A_* and *R_I_*, respectively.

Moreover, we assume that all reactions between species follow the mass-action law. Then for a single ARF and a single Aux/IAA, we have,

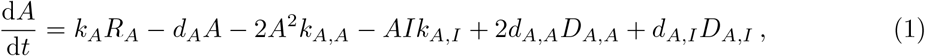

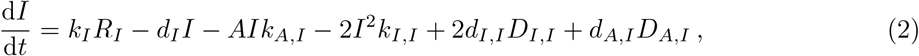

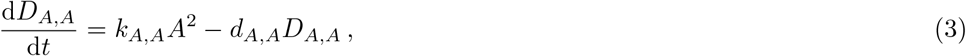

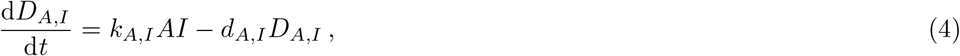

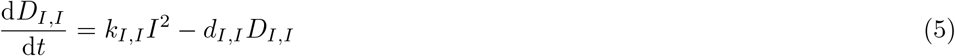

where *k_A,A_, k_A,I_*, and *k_I,I_* are association rates; *d_A,A_, d_A,I_*, and *d_I,I_* are the corresponding dissociation rates; and *k_A_* and *d_A_*, and *k_I_* and *d_I_* are the production and decay rates for ARFs and Aux/IAA proteins, respectively. The production and decay rates for ARFs, that is *k_A_* and *d_A_*, are assumed to be constant.

It is well known that auxin-mediated decay of Aux/IAAs happens via ubiquitination when Aux/IAA is bound to SCF^TIR1^ [23]. This degradation pathway is described more completely by previous models [29, 2, 5], whereby the rate of Aux/IAA decay is dependent not only on the concentration of auxin, but also on the concentration of SCF^TIR1^. For the sake of simplicity, we omit these details, introducing the variable *a* to represent the *effective auxin concentration* such that the decay rates for Aux/IAAs are linearly dependent on *a*,

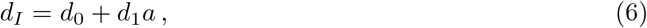

where *d*_0_ is the rate of decay in the absence of auxin, and *d*_1_ is a parameter characterising the auxin dependence of Aux/IAA decay.

To capture transcriptional regulation, we let **g**(*t*) be a vector containing the probabilities that the promoter is bound to a particular monomer/dimer at time *t*. Then, we model the binding of ARFs and Aux/IAAs to promoter regions of DNA according to,

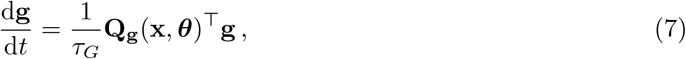

where **Q_g_** is a transition-rate matrix which depends on the models state vector **x** (containing all state variables) and parameter vector (***θ***), and *τ_G_* is some representative timescale that dictates the overall rate of the dynamics, but does not affect the steady state, lim_*t*→∞_**x**(*t*). For fixed **x** and ***θ***, this portion of the model is a conservative Markov model [41].

The state of the promoter at a given time is governed by a Markov model describing dimer and monomer binding to DNA. As described below, these promoter states dictate the transcription rates (affecting mRNA production) and indirectly control the production of ARF and Aux/IAA proteins, as described in Equations (1) and (2). By modifying the transcription rates for each given promoter state, and the resultant mRNA species, we can configure our model such that it describes various transcriptional effects.

This formulation of the transcription module differs from previous approaches such as the models presented by Bintu et al. 2005 [4] and Middleton et al. 2010 [29]. Our formulation describes formation of dimers containing ARFs and Aux/IAAs, and the binding of these dimers with the promoter. By setting binding and transcription rates for particular promoter states, the model can describe competition and cooperativity between regulator complexes. Models can also exhibit transcriptional repression where the binding of a particular complex to the promoter causes a reduction in the transcription rate for a particular gene compared to the basal transcription rate—the transcription rate corresponding solely to activity of the basal transcription machinery and no binding of auxin signalling complexes to the promoter. Moreover, our framework allows the possibility for dimers to form on the promoter itself (though the models presented herein do not have this property).

Using this approach, direct transcriptional repression may be modelled by ensuring that the transcription rate corresponding to the ARF-ARF dimer-bound promoter state is less than the basal transcription rate. Such a scenario where the model includes a repressor ARF is presented in Section 44.2, but for simplificty. Each promoter state corresponds to a different transcription rate for each species of mRNA. There are distinct state variables quantifying mRNA concentrations for each protein in our model, with concentrations denoted by 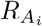 and 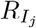, that drive the expression of ARF and Aux/IAA proteins, respectively. Each mRNA concentration, *R_i_*, is governed by an ODE of the form,

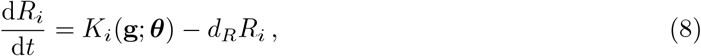

where *d_R_* is the assumed constant decay rate of the mRNA (assumed equal for mRNA concentrations corresponding to any ARF or Aux/IAA species); and *K_i_*(**g, *θ***) is a function which maps the promoter states and parameter vectors to some transcription rate.

Equation (7) describes how the probability that the promoter is in each possible state changes over time. In particular, we denote the components of **g** to be *g_A,A_, g_A,I_* and *g*_0_, representing the probability that promoter is: bound to an ARF-ARF dimer; bound to an ARF-Aux/IAA dimer; and unbound, respectively. The effective transition rate is then,

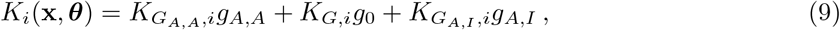

where 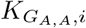 is a constant parameter which quantifies the rate of protein translation when an ARF-ARF homodimer is bound to the promoter, 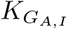 is the transcription rate of gene *i* when an ARF-Aux/IAA dimer is bound to the promoter, and *K_G,i_* is the transcription rate of gene *i* due solely to the basal transcription machinery, when the no ARF-ARF or ARF-Aux/IAA dimers are bound to the promoter due solely to the basal transcription machinery (that is, the basal transcription rate). These variables are model parameters and, as such, are components of the model’s parameter vector, ***θ*** This scenario is shown by the schematic in Figure 4.

**Figure 4.**
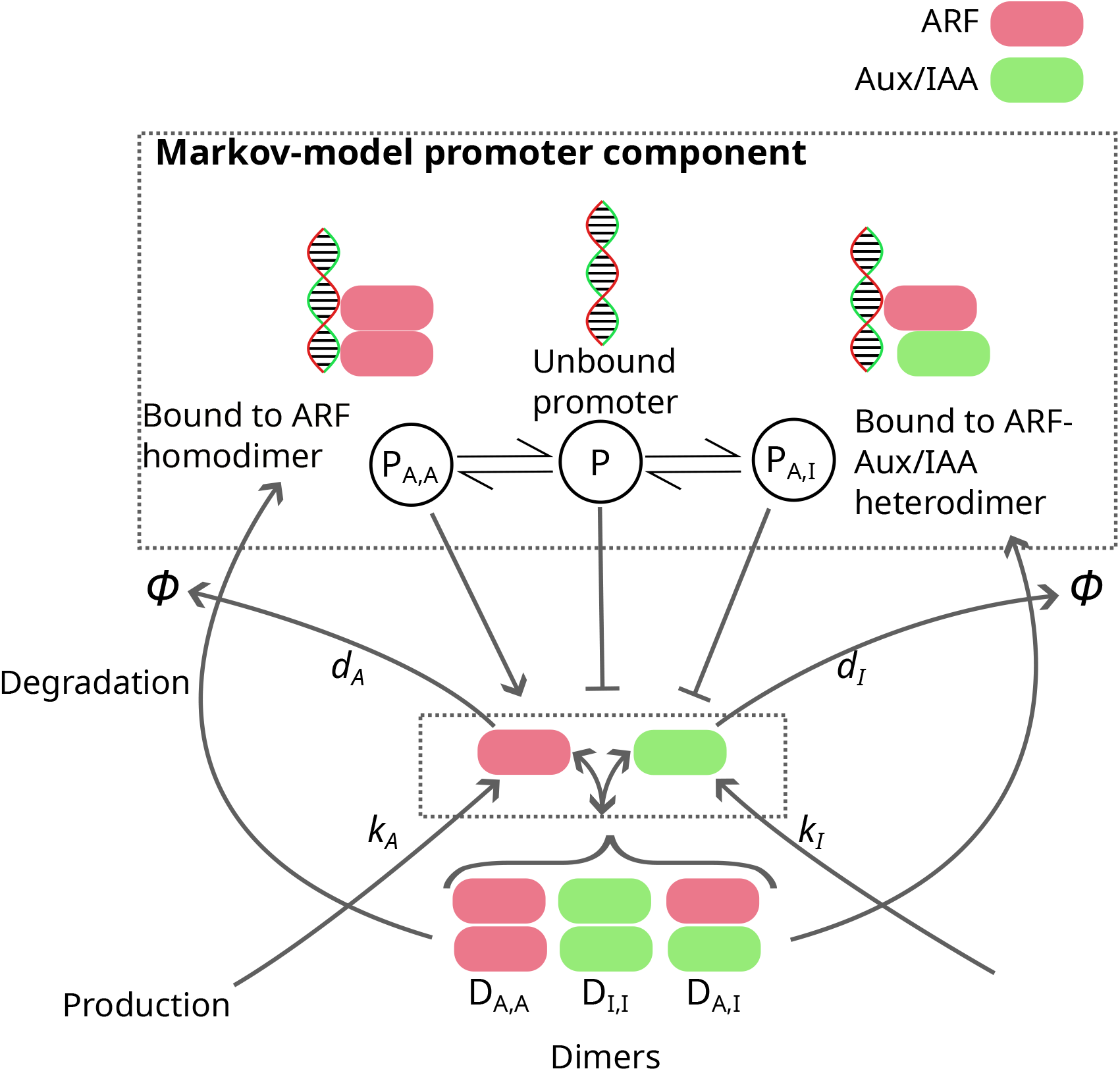
Schematic showing the various components of our NAP modelling framework. A simple, single-ARF, single Aux/IAA NAP is shown for illustrative purposes. The topmost dashed rectangle shows the Markov model describing protein-promoter binding. The probability of the promoter being in each state dictates transcription rates for both ARFs and Aux/IAAs (independently). The lower dashed rectangle shows the production, decay, and dimerisation of ARFs and Aux/IAAs. The concentrations of monomers and dimers affect the transition rates in the Markov model, and hence (indirectly) the transcription rates. Though only a single ARF and Aux/IAA are shown, the same principles can be extended to models with many ARFs and Aux/IAAs. Note that Aux/IAAs lack a DNA-binding domain like that of an ARF, and as such are unable to bind directly to the DNA.

We assume that the promoter-protein dynamics occur rapidly in relation to other parts of the model. Such an assumption is made implicitly in other models, which use probabilistic arguments to define transcription-rate functions [29]. This quasi steady-state assumption allows for the model to be simplified, setting 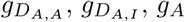 and *g*_0_ by computing the steady-state of Equation (7).

The above-transcription parameters allow a range of activational and repressive effects to be modelled, for every gene represented in the model. For instance, if we consider a single ARF and Aux/IAA, we can omit the subscripts and set 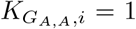 and 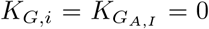 such that transcription of the *i* gene only occurs when an ARF-ARF dimer is bound to the promoter. Consequently, the maximal transcription rate is 1, and as the ARF-ARF dimer concentration, *D_A,A_*, increases, the transcription rate saturates. An example of the relationship between this homodimer concentration, *D_A,A_*, and a given transcription rate is shown in the Supplementary Material (Section S1). Note that when considering multiple ARFs and Aux/IAAs, these formulas will include additional promoter states and state-dependent transcription rates, and so, the relationship between these concentrations and the effective transcription rates may be more complicated.

Another configuration which shows the flexibility of this formulation is where ARF and Aux/IAA transcription is promoted by ARF-ARF homodimers, but repressed by ARF-IAA heterodimers. This is achieved by ensuring a positive basal transition rate, *K_G,i_ >* 0 for each gene *i* (that is, basal transcription of ARF and Aux/IAA). Then we choose 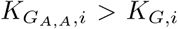 such that ARF-ARF homodimers promote the transcription of both genes, and 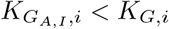, such that ARF-Aux/IAA dimers repress transcription.

In any case, the transcription rate is dependent on the dimer concentrations *D_A,A_* and *D_A,I_*, which are governed by Equations (3) and (4). As *τ_G_* → 0, the timescale of protein-promoter dynamics decreases. For small *τ_G_* we can approximate the system using a *quasi steady-state assumption*,

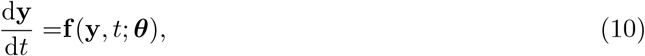

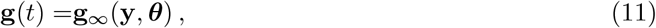

where **f** (*y, t*; *p*) is the vector-valued function describing the evolution of the state variables as in Equation (1)–(5), and the function **g**_∞_ maps these state variables, denotes the equilibrium point of the promoter Markov model corresponding to the chosen parameter values, ***θ***, and the instantaneous state of the model **y** (the concentrations of protein complexes in particular). This equilibrium distribution is determined by finding the non-trivial solution to Equation (7) with elements summing to one, that is, **1**^⊤^**g** = 1. If the protein-promoter dynamics are assumed to happen quickly (compared to other dynamics in the model), the resulting ODE system computed explicitly is “stiff”, meaning that significant computational work is required to simulate the model. This problem is alleviated by the quasi steady-state formulation. Reducing the number of state variables in this way also simplifies the numerical analyses presented in the following sections [41].

### 3.2 General modelling framework for multiple ARFs and Aux/IAAs

We now provide notation for NAP models containing multiple ARFs. Examples II and III below are models of this form, containing two distinct ARFs and a single Aux/IAA species. Because it is simple to extend this modelling framework to an arbitrary number of ARFs and Aux/IAAs, we provide the notation and software tools required to define and analyse such models. However, we do not consider any modles with multiple Aux/IAA in this article. Nevertheless, we introduce the necessary notation.

We use *A*_1_, *A*_2_, …, *A_N_* to denote the concentration of *N* different ARF species. Similarly, we notate concentrations of the various Aux/IAA species as *I, I*_2_, …, *I_M_* where *M* is the number of Aux/IAA species represented in the model. Then, we denote a dimer of the *i*^th^ ARF transcript with the *j*^th^ ARF transcript as 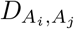. Likewise, 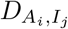 denotes the concentration of dimers formed from ARF *i* and Aux/IAA *j*, and 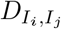 denotes the concentration of dimers formed from Aux/IAA *i* and Aux/IAA *j*. As before, we assume that the order of the indices of the dimer are inconsequential.

Using this notation we can describe the dynamics of NAP with multiple ARF and/or Aux/IAA species under the modelling assumptions listed above. For example, for a model including two distinct ARF species, but only a single Aux/IAA species, we have,

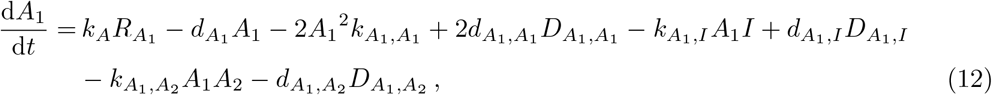

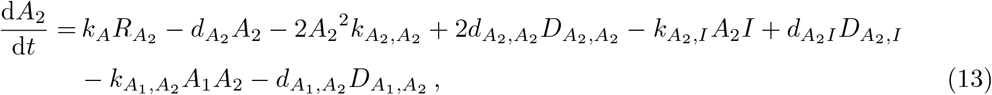

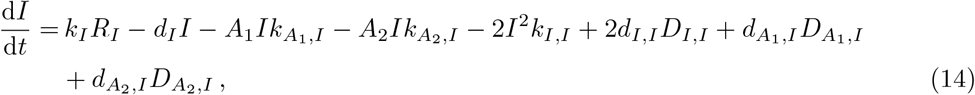

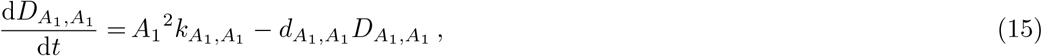

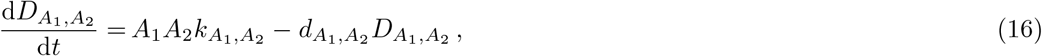

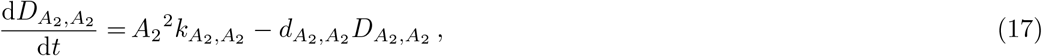

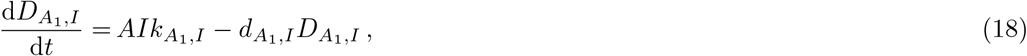

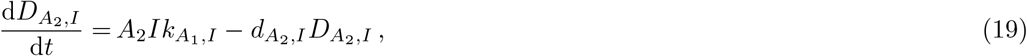

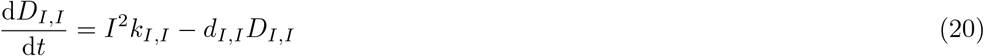

As in the single-ARF, single-Aux/IAA model, the concentration of each species of mRNA is governed by Equation (8), and the corresponding effective transcription rates computing via the same Markov-model formulation as in Equations (7) and (9). Moreover, we again simplify this model by omitting the monomer-bound promoter states and using a quasi steady-state assumption (as in Equations (10)–(11)), where protein-promoter binding events are assumed to occur much more rapidly than other processes in the model. The resulting quasi steady-state of the promoter Markov model then determines transcription rates similarly to Equation (9), but with additional terms for the additional promoter states.

For the examples provided in this section, the model is further simplified by the assumption that lone ARFs can not be bound directly to the promoter, and that dimerisation of a promoter-bound monomer is not possible. In this case, the Markov chain described by Equation (7) has a simple “hub-and-spoke” topology (where each promoter-bound state is directly connected only to the unbound state), such that the steady-state probabilities are given by,

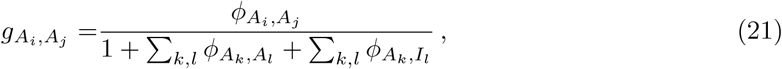

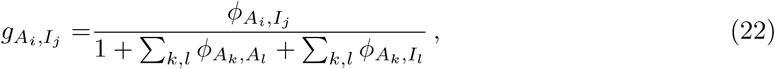

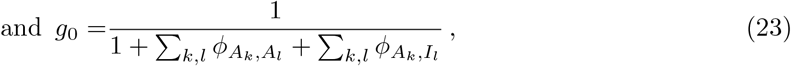

where 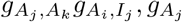 and *g*_0_ denote the probability that the promoter is in each of the given states, and 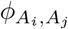 and 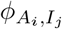 denote the ratio of association and dissociation rates for ARF-dimer-bound promoter states, and ARF-Aux/IAA-dimer-bound states, respectively. Note how this quasi steady-state assumption effectively reduces the number of necessary parameters, as the above steady-states may be expressed simply in terms of ratios association and dissociation rates.

Accordingly, for a model with *N* ARF species and *M* Aux/IAA species, we may write the effective transcription rate by combining the state-dependent transcription rates for each of the promoter states (with occupation probabilities denoted, as above, by 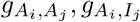 and *g*_0_),

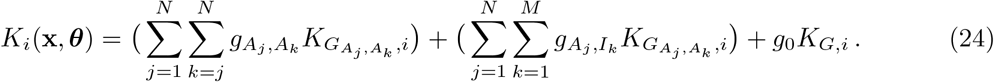

Note that this formulation could be extended as to also include monomer-bound promoter states. However, this is not necessary for the work presented here.

Whilst such models may be cumbersome to present in full (especially as the number of signalling components increases), the implementation of such models in computer code is straightforward. Our software package `auxin signalling_models`(available as a GitHub repository) allows for the generation such models, with a user-defined number of ARF and Aux/IAA species. No matter the precise configuration of ARF and Aux/IAA transcripts, we can make use of the assumption of a quasi steady-state, as the promoter component in each model always has a unique steady state provided the underlying network is connected [41].

In the following section, we put our framework to use and explore three example models of increasing complexity.

## 4 Applications of NAP modelling framework

We start with a single-ARF, single-Aux/IAA model describing ARF5 and IAA12, a well-studied ARF-Aux/IAA pair. Computationally and experimental investigations have suggested that this module can exhibit bistability [21]. We demonstrate that bistability is also possible under our model, validating our approach. Following this example, in Section 44.2, we add a repressor ARF to this model and demonstrate how controlling the expression level of this additional ARF can modulate the module’s behaviour. Finally, in Section 44.3, we construct a module consisting of two activator ARFs and explore its response under step-change and sinusoidal auxin signals. A diagram of each example is shown in Figure 5.

**Figure 5.**
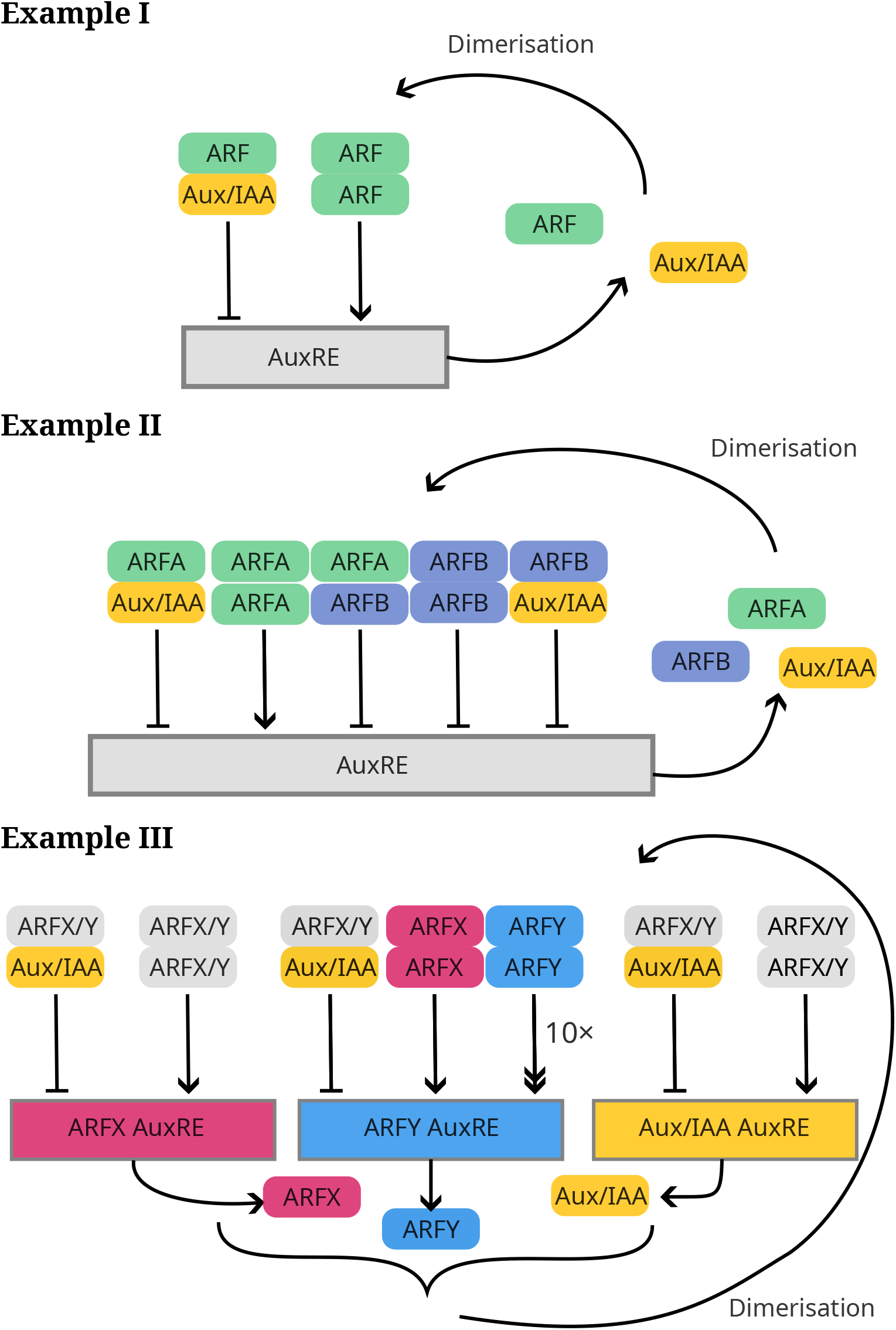
Diagrams of the example NAP subnetworks explored in this article. **Example I**: A single-ARF, single-Aux/IAA model where transcription (of ARF and Aux/IAA genes) is promoted by ARF homodimers, but repressed by ARF-Aux/IAA heterodimers. **Example II**: the previous model but with an additional repressor ARF with mRNA transcribed at a constant rate unless the promoter is bound to a complex containing Aux/IAA (not pictured). **Example III**: A double-ARF feed-forward motif in which two class-A ARFs (ARFX and ARFY) promote each other and where one ARF, ARFY, promotes itself particularly strongly.

### 4.1 Example I: single-ARF model

We begin with a single-ARF, single-Aux/IAA example. This example may be expressed in the general notation laid out in the section above, but we choose to omit unnecessary subscripts for brevity. One well-studied combination of ARF signalling pathway components is ARF5/MONOPTEROS (MP) together with Aux/IAA12/BODENLOS (BDL). Previous work includes a model where MP promotes transcription of MP and BDL, but where MP and BDL work cooperatively to repress the same genes [21]. In [21], the authors suggest that the mechanism for this cooperativity is the formation of MP-BDL dimers which, when bound to the promoter, repress both MP and BDL transcription. In this section, we use the general model introduced above to model this specific network. A diagram of this network motif is shown in Figure 5. We use the same governing equations as before (Equations (1)–(5)) to describe the production and decay of monomers, and the association and dissociation of dimers.

To capture the promotion and co-repression dynamics described above, we set the state-dependent transcription rates to be,

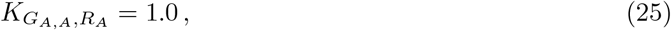

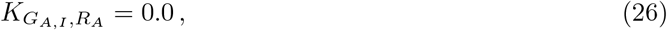

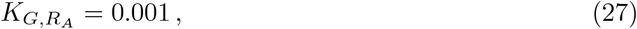

and similarly, those for BDL to be,

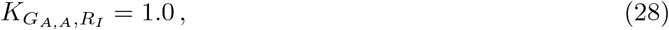

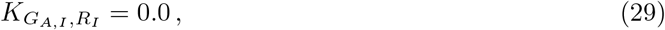

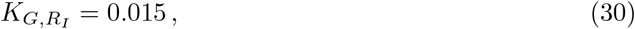

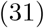

where the first subscript (a dimer) corresponds to the complex bound to the promoter region, and the second subscript denotes the type of mRNA being produced—MP (*A*) or BDL (*I*); the basal transcription rates where no compound is bound to the promoter are denoted by *K_G,A_* and *K_G,I_* for MP and BDL, respectively.

From these state-dependent transcription rates, we see that an increase in BDL-MP dimers represses transcription of MP and BDL because this (rapidly) increases the probability that an MP-BDL dimer is bound to the DNA, and 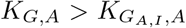 and 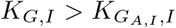. Likewise, we see that an increase in ARF-ARF dimers effects an increase in transcription of the same genes because 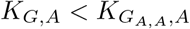 and 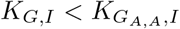.

It has been demonstrated *in vitro* for some ARFs that dimer-promoter binding is much more stable than monomer-promoter binding [14], so we make the simplifying assumption that no ARF monomer can bind to DNA. This has the effect of simplifying the promoter component of the model (that is, the Markov model characterised by **Q_g_** in Equation (7)) such that its graph representation has a simple topology where each dimer-bound promoter state is connected only to the unbound state. In such a case, only the ratios of association and dissociation rates (binding thresholds) are important, and it is simple to write down the equilibrium distribution which takes a similar form to the transcription rates presented in [29].

By default, all state variables are initialised with *A*(0) = 1.0 and all other state variables set to 0.1. The system is then equilibrated by simulating with a fixed effective auxin concentration, *a* = 0.1, until *t* = 48 hours. A full table of model parameters is provided in the Supplementary Material (Section S2)

We performed a manual exploration of parameters and identified a regime where this model exhibits multistability, as shown in Figure 6. For this particular parameter set, we see three branches of equilibria in Panel **a**: stable equilibria on the topmost branch, a connected branch of unstable equilibria and a second branch of equilibria where the total Aux/IAA concentration is much lower, especially for *d_I_ >* 0.01. The behaviour of the model around these equilibria is explored further in Panels **a** and **b**, where we see the effects on an apparent unstable limit cycle.

**Figure 6.**
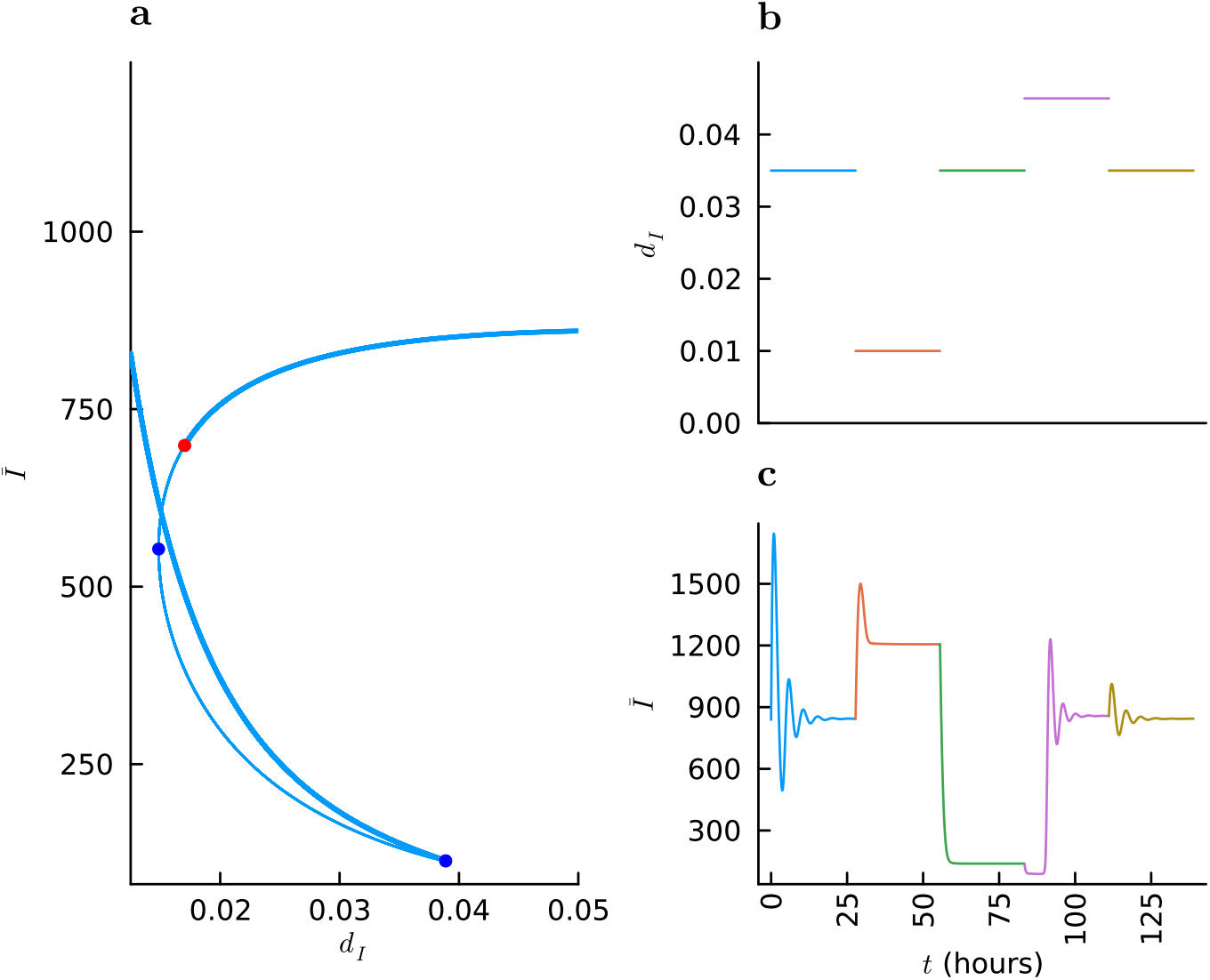
Multistability in the NAP can lead to hysteretic responses for certain auxin concentrations, as shown by the total (bound and unbound) Aux/IAA concentration, *Ī* = *I* + 2*D_I,I_* + *D_A,I_*. Three different auxin concentrations are applied to the model, and we see that a drop in auxin sends the model to a sole stable branch with high *Ī* (shown in Panel **a**), and the application high auxin con-centration returns the network to the original branch. For auxin intermediate auxin concentrations, however, the system may converge to one of two stable equilibria. Panel **a**: bifurcation diagram showing possible steady states; stable steady states are marked by the thicker blue line and unstable steady states by a thinner line; blue circles show the location of bifurcation points and the red circle denotes a Hopf bifurcation. Panel **b**: a piecewise constant auxin input signal. Panel **c**: the model’s response to the above auxin signal in *Ī*.

Note that the bifurcation parameter, *d_I_*, encodes both auxin-dependent and auxin-independent decay rates such that as the auxin concentration in the cell increases, so too does the decay rate, *d_I_*. Therefore, the behaviour shown in Figure 6 may be exhibited for any form of *d_I_* (such as a hill function), so long as the decay rate increases with *a*, and suitable parameters and auxin concentrations are provided. For sufficiently small *d_I_*, where little Aux/IAA decay occurs, one branch becomes unstable, and only one stable equilibrium remains. In general, we may expect that multistability is found only for specific ranges of auxin concentration and that for sufficiently large auxin concentrations, there is a single monostable regime where the total Aux/IAA concentration, *Ī*, is small and transcription rates (for *A* and *I*) approach their maximal values.

This interesting dynamic behaviour (namely multistability and hysteresis) arises naturally from the proposed binding kinetics and the transcription rates under different promoter states as introduced above, albeit with particular choices in parameter values. The model presented by Lau et al. [21], of the same MP-BDL subnetwork, exhibits bistability, so finding similar behaviour in our model (through manual parameter-space exploration) is not unexpected. Compared to the full NAP found in *Arabidopsis thaliana* (consisting of all the proteins coded for in the genome), the MP-BDL subnetwork described by Lau et al. [21] and this section is very simple, containing only a single ARF and Aux/IAA species.

The qualitative behaviour of such models is associated with the presence of positive and negative feedback loops—the former being a necessary condition for the existence of multistability, and the latter being necessary for sustained oscillatory behaviour [43]. Unlike the simulations presented in [21], however, our model exhibits damped oscillatory behaviour, as shown in Figure 6.

These mathematical insights and *in silico* observations align with experimental observations showing that bistability in MP-BDL dynamics plays a key role in embryogenesis and cell-fate determination in *Arabidopsis thaliana* [21, 35]. Oscillatory behaviour is known to be important to root development in *Arabidopsis thaliana* [6], but, though plausible, similar oscillations have not been observed in the MP-BDL subnetwork [19].

This single-ARF, single-Aux/IAA model serves as an example of the complex dynamics can emerge even from simple subnetworks of the NAP with few components, exhibiting bistability, oscillatory transient behaviour smooth graded responses to changes in auxin concentration. All these classes of response are likely to co-exist within the same tissue, but with different subsets of genes being regulated. For example, within the root meristem data shown earlier there should be bistable genes associated with cell identity [31], graded responses in cell elongation proportionate to auxin concentration that control root growth and gravitropism [2] and oscillations that control lateral root formation [10]. These simulations reveal that a single ARF-Aux/IAA model is capable of producing these outcomes, with different outcomes dependent on parameters. Based on the fact that we know all these responses co-exist within a single tissue, this highlights the need for including multiple ARF-Aux/IAA subnetworks each with different parameters and feedbacks.

Given the simplicity of this example, we may expect that similar networks play other important roles in *Arabidopsis thaliana* and other angiosperms, though any examples may be complicated by other NAP interactions and so be difficult to isolate experimentally. In the next section, we move beyond this single-ARF, single-Aux/IAA setting and further utilise flexibility modelling framework by exploring the action of a repressor ARF when added to the same MP-BDL system.

### 4.2 Example II: promoter and repressor ARF model

Using the general model outlined in the previous section, we can construct models with additional ARF species. Here, we gradually add some complexity to NAP subnetwork under investigation, adding a class-B repressor ARF to the network described above. A schematic for this example model is shown in Figure 5. In Figure 3(**a**), we saw that some class-B repressor ARFs showed a noticeable transcriptional response to auxin treatment. One such example is ARF4, which showed a steady increase in expression as the duration of auxin treatment increases (Fig. 3 **a**). Previously, it has been shown experimentally that ARF4 interacts competing with MP for BDL binding sites [52].

We proceed by including a repressor ARF, which we refer to as ARFB, in the model. Then there are two separate ARF concentration variables, *A_A_* and *A_B_* which represent concentrations of the activator ARF, MP, and repressor ARF, respectively. We also denote the various dimers formed by these ARFs and Aux/IAAs using the variables, 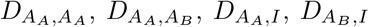, and *D_I,I_*, as introduced in the Section 3.2. We assume both these ARFs, and Aux/IAA, are induced by auxin, and, where possible, we use the same parameters as in the previous example.

Following the previous section, we then have additional state variables to represent various dimer concentrations: 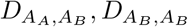, and 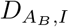. We assume that these two ARF species behave identically, except the transcription rate of the repressor ARF is an equal, positive value in all cases except where an Aux/IAA is present in the promoter-bound complex. That is, by setting,

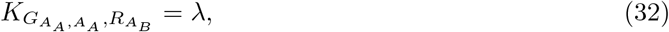

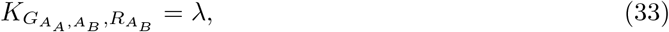

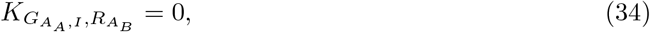

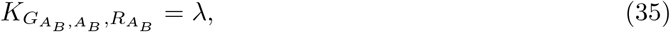

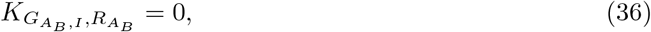

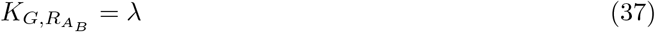

where *λ* is the aforementioned transcription rate.

The repressive effect of the newly-added ARF is achieved by assuming no transcription of ARFA 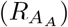 or BDL (*R_I_*) whenever the promoter is bound to a dimer containing ARFB. That is,

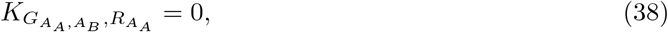

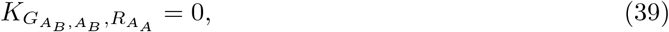

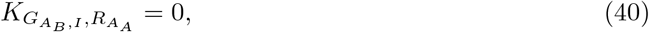

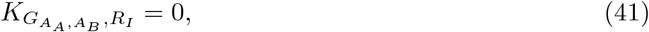

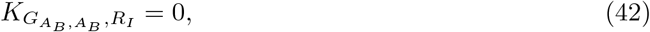

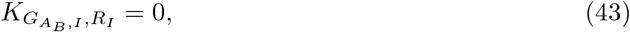

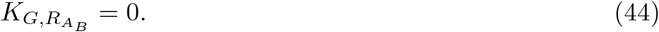

In this way, an increased ARFB concentration (say from *λ* being increased) leads to reduced transcription of MP 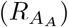 and BDL (*R_I_*), as expected. Increasing *A_B_* also leads to increased formation of 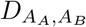 dimers, reducing *A_A_* and hence 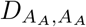. Again, a full table of model parameter values is provided in the Supplementary Material.

In Figure 7, we see that this system has a bifurcation point and bistability provided the ARFB transcription rate (in the case that no dimer containing Aux/IAA is bound to the promoter), *λ*, is not too large. In other words, we see that an increase in repressor-ARF activity can eliminate bistability.

**Figure 7.**
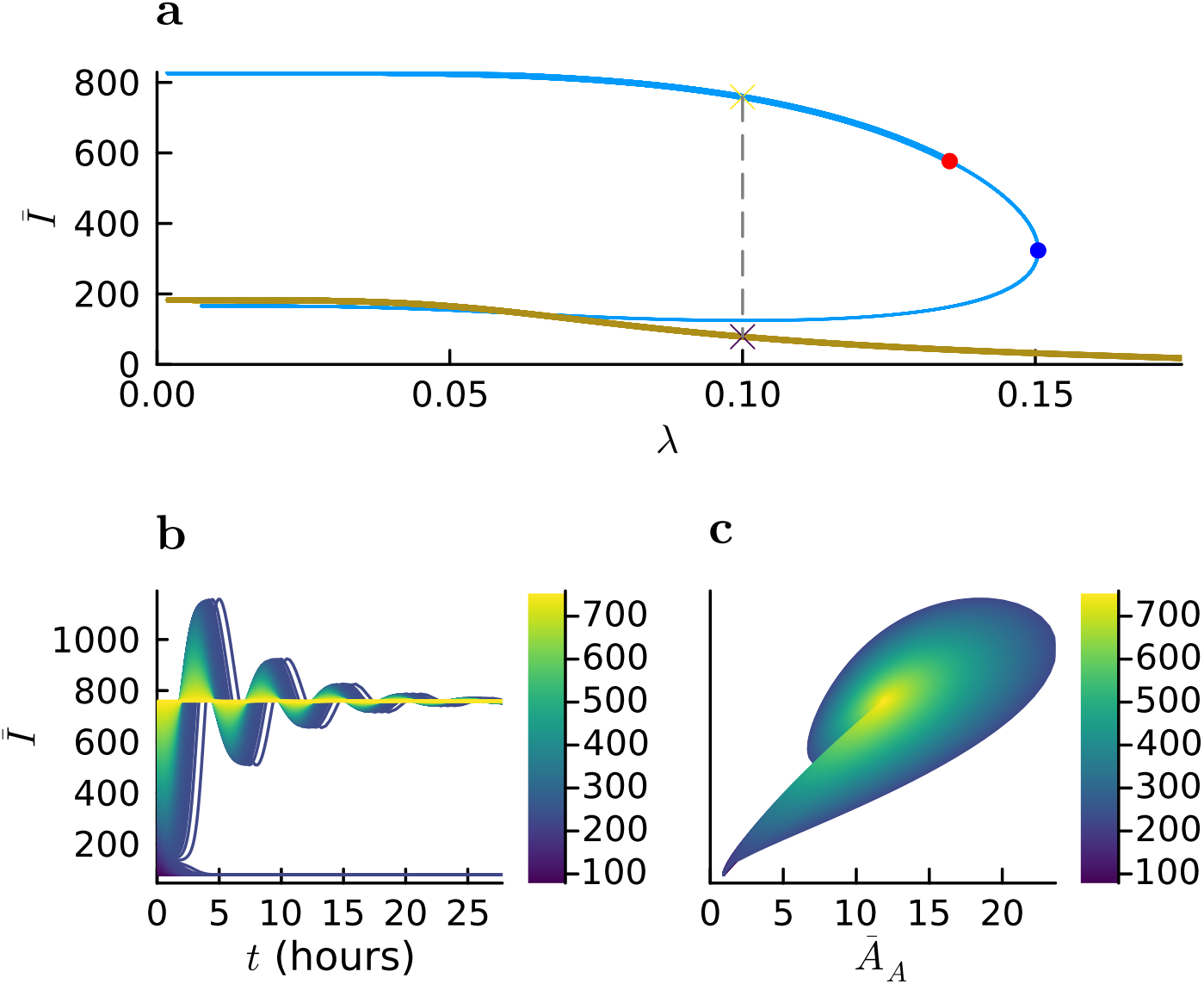
Provided repressor-ARF activity is not excessive, the MP-BDL model (augmented with a repressor ARF) can be multistable. The total BDL concentration (both bound and unbound), *Ī* is shown on the y-axis. **a** shows a bifurcation diagram with a bifurcation point and bistability between two stable branches (shown in thick blue and brown lines) for various values of the transcription-rate *λ*. The thin blue line shows a branch of unstable equilibria. A subcritical Hopf bifurcation is displayed in red, and a branch point is shown in blue. **b**: simulations under the same parameter set using various initial conditions selected along the dashed line segment shown in Panel **a.** Panel **c**: phase-plane plot of the same simulated trajectories, showing total BDL concentration (bound and unbound), *Ī*, vs. total MP concentration, *Ā*.

The above example shows how action of a repressor ARF might be described mathematically, and shows a possible mechanism for the deactivation of any given signalling motif in the NAP. A repressor ARF (ARFB in this example) could be transcriptionally regulated in many ways (by other parts of the NAP, or by other signalling networks entirely) modifying the transcription rate, *λ*. In this way, certain subnetworks (operating as logic units such as binary switches), might be be activated or deactivated according to environmental and developmental cues.

Example I showed that a single-ARF, single-Aux/IAA network could act as either a bistable or monostable circuit depending on parameters. By including a repressor ARF, we observe that there are a greater number of scenarios in which the network output can switch between bistable and monostable regimes. This switch can happen independently of parameters present in Example I (including auxin concentration) and can be obtained solely by altering regulation of the repressor ARF. This capability would mean that external cues regulating a repressor ARF could modulate the outcome of a subnetwork containing this component to provide a response distinct from other subnetworks without that ARF.

### 4.3 Example III: coupled activator ARFs

Where the previous section explored a simple motif consisting of a single activator and single repressor ARF, this section explores a theoretical interaction between two hypothetical activator ARFs. In Section 2, we observed that many ARFs are not strongly transcriptionally regulated by the first time-point, 30-minutes after exposure to auxin. One explanation for this is that their auxin-responsive transcriptional response occurs via the regulation of an intermediate gene. That is, the gene exhibiting a slow regulatory response to auxin may not be a *primary target* of auxin, but may instead be regulated by upstream ARFs that are themselves primary targets. Such slower transcriptional responses are common, as shown in Figures 2 and 3.

Transcriptional activation is known to occur between ARFs [16], and may explain why we observed that certain ARFs do not respond strongly until 60–120 minutes after auxin treatment. To explore this idea further, we present a subnetwork in which two activator ARFs, ARFX and ARFY, exhibit different responses to the same auxin stimulus: one ARF (ARFX), being more sensitive and responding quicker, whilst the second (ARFY) primarily responding to activation by the first. We then show how tuning particular parameters in this subnetwork can lead to different response times in the second, downstream ARF, but not the first. We suggest that similar regulatory motifs may explain the differences in response times exhibited in Figure 3 [16].

We refer to the two hypothetical self-activating, class-A ARF species as ARFX and ARFY, with concentrations denoted *A_X_* and *A_Y_*, respectively.

We configure the model such that ARFY is induced by ARFX, and so for certain parameter regimes, upregulation of ARFY in response to an increased auxin concentration occurs much slower than for ARFX. This situation is shown in Figure 5. In particular, we focus on the rates governing dimerisation of ARFY with itself, ARFX and Aux/IAA: the association rates, 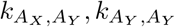 and 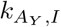; and the dissociation rates, 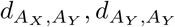 and 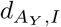. We show that the speed of ARFY transcriptional response can be controlled by simultaneously scaling these parameters.

Whilst we modify the ARFY dimerisation rates, the transcription rates for ARFX, ARFY and Aux/IAA stay the same throughout this section. For ARFX, we assume that transcription occurs in all promoter states, but less so when an Aux/IAA is bound to the promoter region, by setting 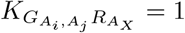 and 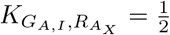, for either ARF, that is *A* = *A_X_* or *A* = *A_Y_* with *i, j* ∈ *{X, Y*}. Hence, we expect the ARFX transcription rate to rapidly increase in the presence of auxin due to an increase in Aux/IAA decay, which causes a relative increase in the amount of ARF-ARF dimers present (and as a result, an increased probability that an ARFX-ARFX dimer is bound to the promoter).

For Aux/IAA, we assume transcription happens only if no dimer containing an Aux/IAA is bound to the promoter, and we set our transcription rates to be: 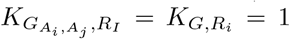 and 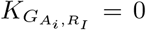. Using these constraints, our model describes a network where ARF and Aux/IAA transcription is promoted by both ARF species, and ARF and Aux/IAA transcription is inhibited by Aux/IAA. A full table of model parameters is presented in the Supplemental Material.

Then, for ARFY we suppose that the transcription only occurs when the promoter is bound to an ARF-ARF dimer, that is, 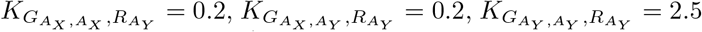, and all other promoter-dependent ARFY transition rates (including the basal transcription rate) are zero: 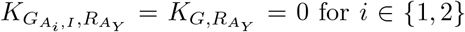 for *i* ∈ *{*1, 2}. In this way, ARFY self-promotes and is promoted (less strongly) by ARFX, and repressed by Aux/IAA. As for previous sections, complete tables of model parameters and initial conditions may be found in the Supplementary Material.

A maximal ARFY transcription rate is obtained when the probability of an ARFY-ARFY dimer being bound to the promoter is 1, that is, when there is a large amount of ARFY-ARFY dimers present (both in absolute terms, and relatively when compared to the other compounds). However, we also expect there to be low amounts of ARFY present in a low-auxin environment, because the basal transcription rate is 0. Therefore, we may expect a response to high-auxin concentration starting with a small rapid increase as ARFX-ARFX dimers bind to the promoter (increasing ARFX and ARFY transcription). Following this, depending on the speed of ARFY dimerisation, we may see a slower sustained increase as more ARFY is produced and more ARFY-ARFY dimers form and bind to the promoter region. A full table of model parameters is provided in the Supplementary Material.

Under these assumptions, certain functional characteristics of this network may be modulated by modifying association/dissociation rates of particular dimers. In particular, we fix the ratio of association and dissociation rates for ARFY dimerisation with itself, other ARFs, and Aux/IAA, scaling these rates together. That is, we introduce a scaling factor *κ*, and update the association/dissociation rates for ARFY-ARFY dimerisation as follows,

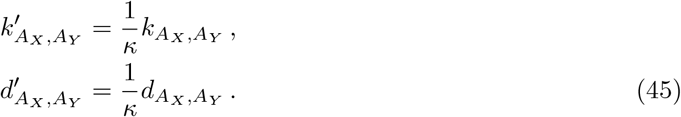

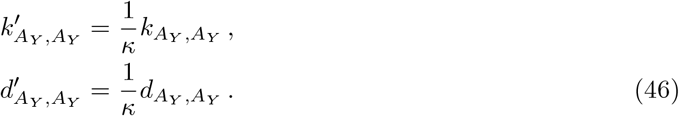

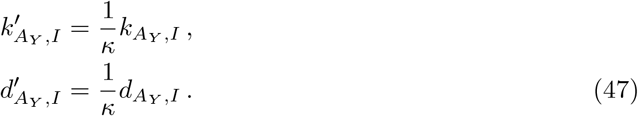

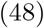

In this way, increasing *κ* slows down the dimerisation of ARFY with other compounds (including itself), without affecting the model’s equilibrium concentrations (as demonstrated in the Supplementary Material B). Only the transient behaviour of the model is affected, causing changes in the time-taken for ARFX and ARFY to respond to auxin stimulus, as demonstrated in the following section.

Broadly speaking, as the effective auxin concentration, *a*, increases, the total Aux/IAA concentration, 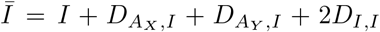 (including both bound and unbound forms) decreases, though for some auxin concentrations the model exhibits multi-stability, as shown by the bifurcation diagram in Figures 8 (panels **a** and **c**). Aside from affecting steady-states, parameters have a large impact on the transient response of the model to different auxin stimuli. We proceed to explore how dimer association and dissociation rates affect the time taken to produce transcriptional responses to step-changes in auxin concentration. Then, we consider how these same parameters affect characteristics of the models’ response to oscillatory auxin stimuli.

**Figure 8.**
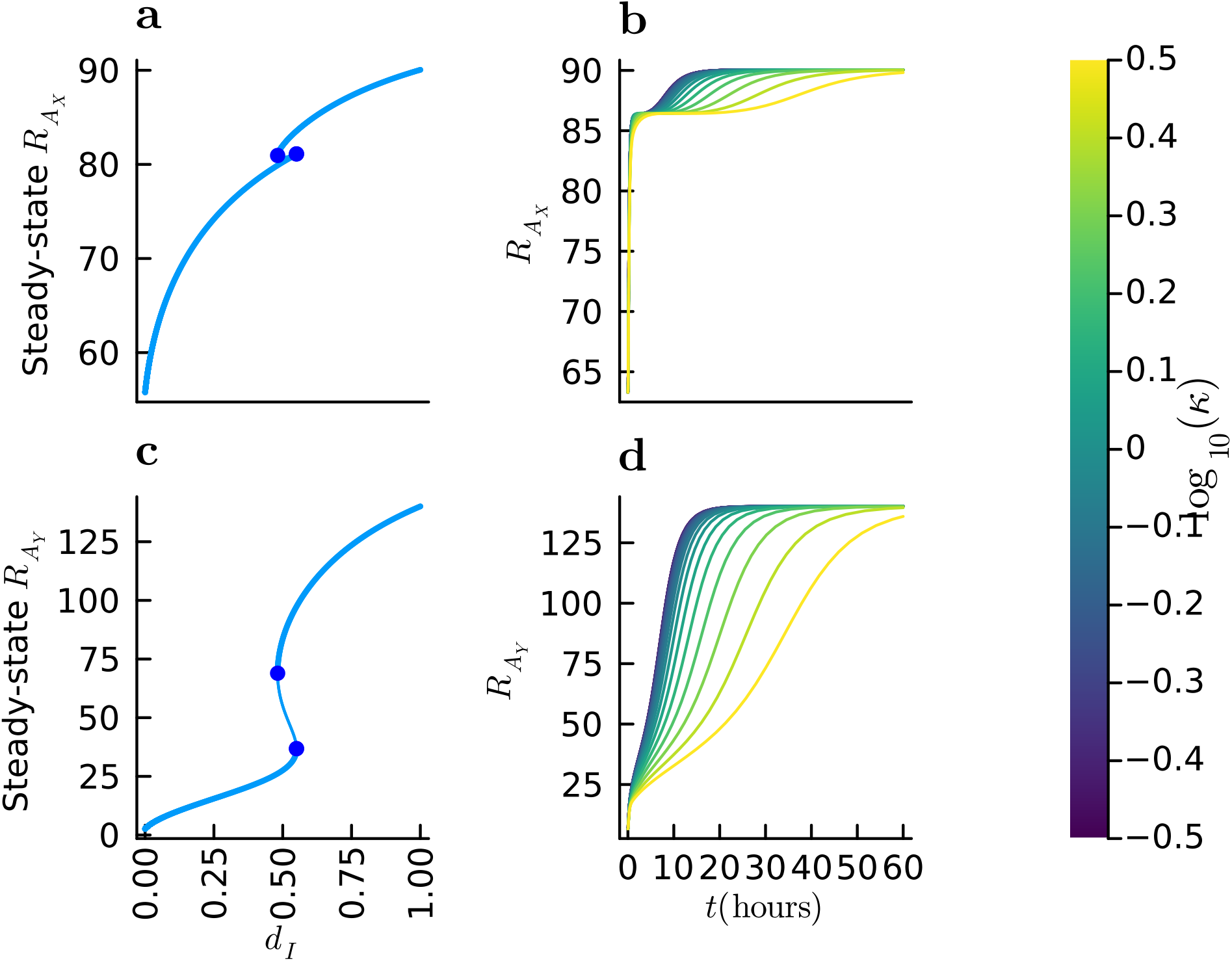
Transcriptional response to auxin treatment in 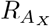 and 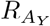 shown for various values of *κ* found by equilibrating the model under low-auxin conditions and simulating under high auxin. **a**: bifurcation diagram showing steady states, denoted by blue circles, as the auxin concencentration, *a*, varies. **b**: a selection of ARFX-mRNA trajectories obtained by simulating the model with different values of *κ*. Panels **c** and **d**: show similar plots for ARFY mRNA, that is 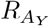. For high *κ*, ARFY dimerisation occurs slowly, causing a delayed response. Whereas, the speed of transcriptional response for ARFX varies little under different values of *κ*.

#### 4.3.1 Auxin response time

Figure 8 shows how larger values of *κ* introduces a delay to the auxin-induced upregulation of ARFY under this model. Whilst the transcriptional response of ARFX varies little under these changes in *κ*, the time taken for the upregulation of ARFY is controlled by this parameter.

#### 4.3.2 Response to sinusoidal auxin concentration

Temporally oscillating auxin signals and fluctuations are found in plants with various periods— the root clock [30], for example, or auxin rhythmic centrifugal waves in the shoot [15]. It is also known that there is interplay between the nuclear NAP and cell-level auxin transport processes (such as transport via PIN and AUX/LAX proteins). Accordingly, we expand our attention beyond constant auxin concentrations and consider the application of a sinusoidal auxin stimulus. We model a representative oscillatory signal by fixing our model’s intracellular auxin concentration to a sinusoidal function, 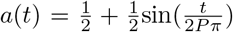, where *P* is the period of the oscillation. An example simulation of the model is shown in Figure 9.

**Figure 9.**
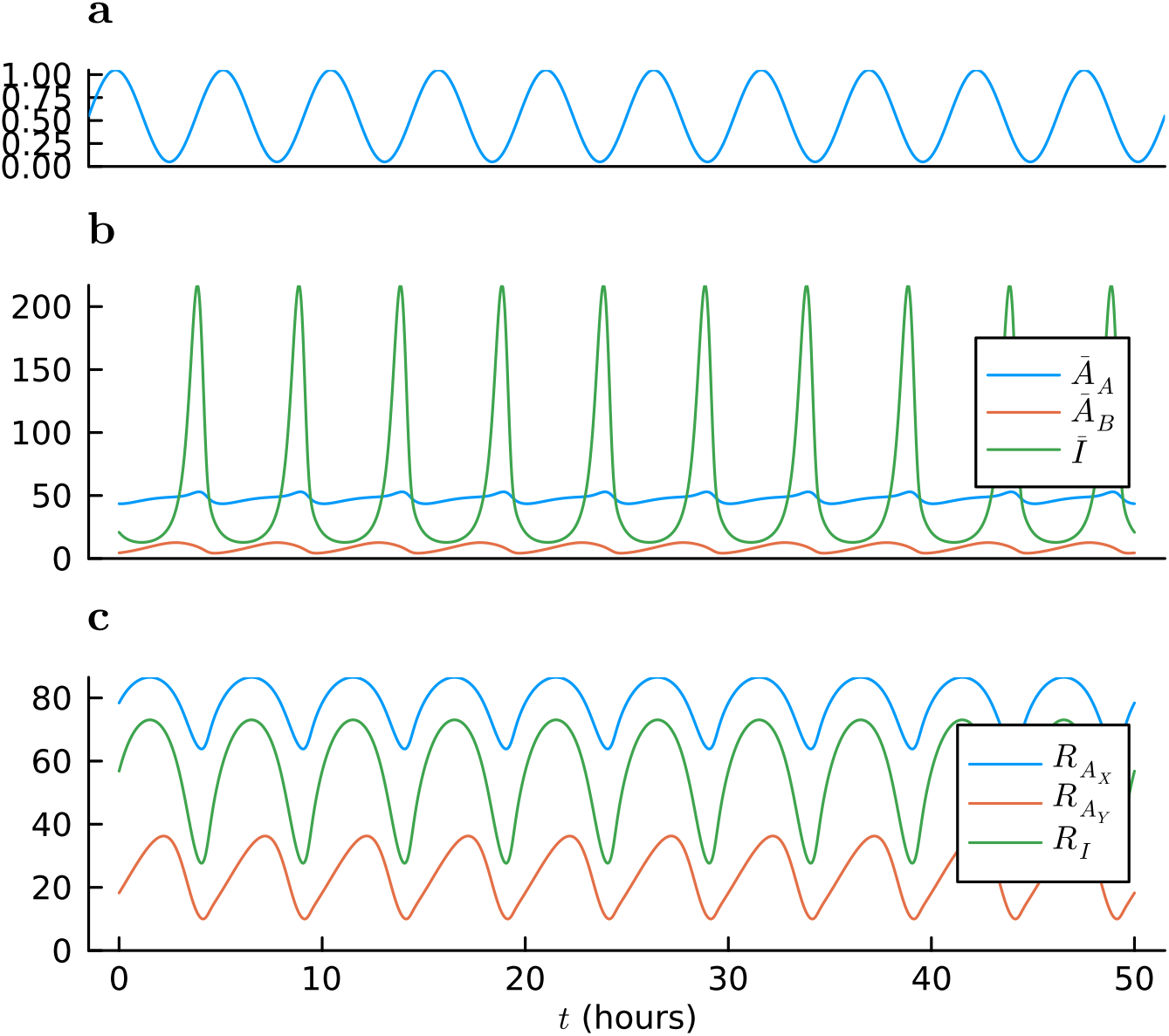
Output of the simulated model when *κ* = 100 and *P* = 5hours. The Aux/IAA concentration rapidly increases and then decreases during the portion of the signal where auxin is low. This, through its binding with ARFX and ARFY, prompts a transcriptional response for ARFX, ARFY and Aux/IAA. However, these transcriptional responses have different characteristics: 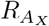 has a roughly steady-state expression when compared to 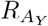 which exhibits relatively larger oscillations. Panel **a** the auxin signal with period *P* = 5 hours. Panel **b**: total protein concentration of all complexes containing ARFA, ARFB and Aux/IAA (*Ā_A_, Ā_B_, Ī*, respectively). Panel **c**: mRNA concentrations for the genes coding for ARFX, ARFY and Aux/IAA (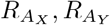, and *R_I_*, respectively).

To characterise the resulting mRNA output we compute,

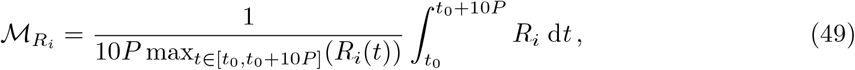

where *P* is the period of the auxin input and *R* is some mRNA concentration, and where *R_i_* is the time-course of interest, being an mRNA concentration such as 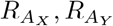, or *R_I_*. In words, 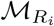 indicates whether the resulting signal is oscillatory or not. For a constant signal, we would have 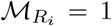, and for a signal consisting of short, rapid spikes we would have 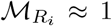. Since we wish to characterise the long-term behaviour of the model at its limit cycle, so we compute 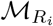 by simulating the model until *t* = 1, 000*P*, and computing 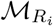 using *t*_0_ = 990*P*.

Figure 11 demonstrates how the functional output of the model depends on *κ* and the period of the input signal (auxin concentration). Certain frequencies of auxin input lead to qualitatively different ARFY transcriptional responses provided a suitable value of *κ*. This demonstrates how the dimerisation dynamics of ARFs in the model may give rise to different temporal expression profiles for target genes in response to non-constant auxin signals.

A heatmap showing 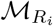 for both ARFX and ARFY, and various (*κ, P*) for both ARFX and ARFY is shown in Figure 11. Little difference is found in 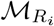 for ARFY across our selection of (*κ, P*) pairs. However, there are noticeable differences in the behaviour of 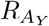 as shown in more detail in Figures 10, where 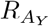 is plotted under numerous simulations using various (*κ, P*) values. Here, we see that the maximal ARFY response in each oscillation is greater for smaller values of *κ* and that the position of these maxima is shifted when compared to the high-*κ* trajectories.

**Figure 10.**
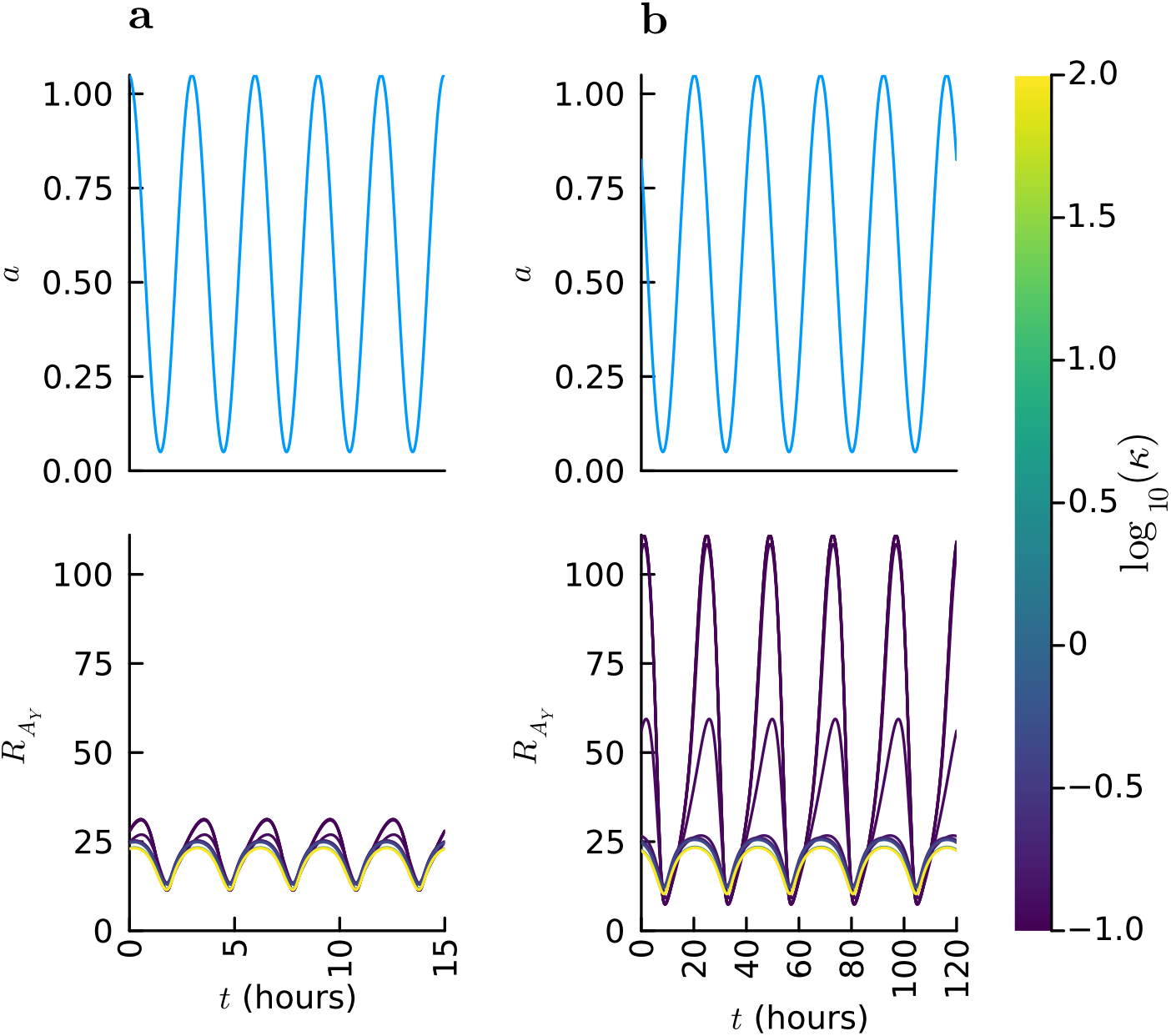
Normalised trajectories for two different auxin-input periods. Column **a**: simulation of a signal with period *P* = 3 hours. Column **b**: simulation of a signal with period *P* = 24 hours. Note that not enough periods have been simulated for the model to reach steady state in some cases. Here we see that the relative size of oscillations depends on *κ*, with larger *κ* leading to relatively small oscillations, and smaller *κ* leading to relatively large oscillations. This effect is particularly noticeable for lower-frequency stimuli (that is, for larger *P*).

We summarise the mRNA concentrations for both ARFs in Figure 11. In this example, the rate of ARFY dimerisation controls the sensitivity of the model output to *P*, the period of the auxin input. The output signal is most sensitivity (as quantified by Equation (49)) to changes in the auxin-signal period when dimerisation is rapid (that is, for small *κ*). Notably, the expression profiles of ARFX are much more uniform than those of ARFY across different values of *κ*.

**Figure 1.**
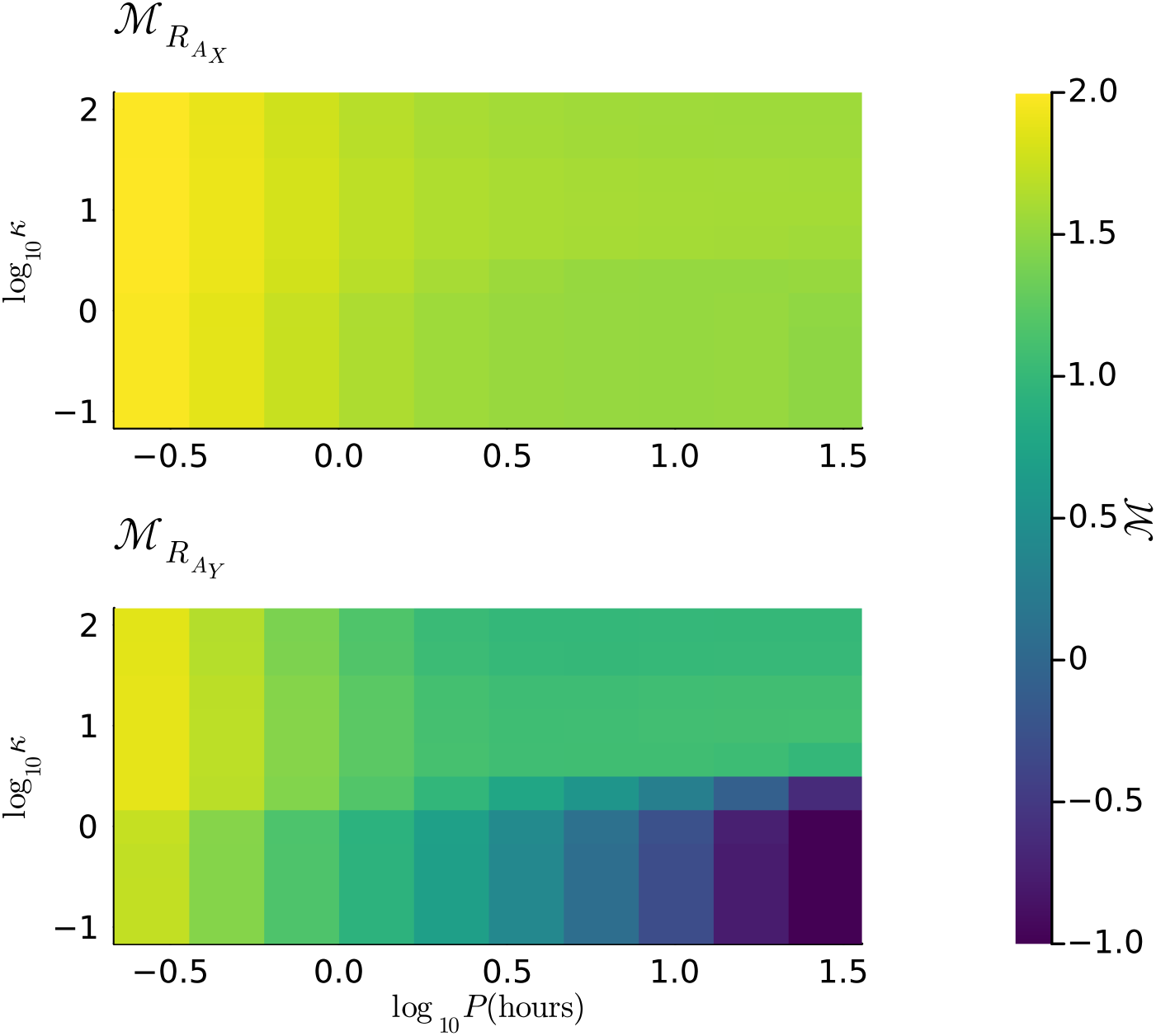
A heatmap showing 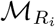 for various combinations of *κ* and *P* for both ARFX mRNA and ARFY mRNA, with *κ* ≈ 1 signifying constant expression levels. The ARFY transcriptional response to auxin signals with different inputs is largely dependent on the timescale of ARFY dimerisation (especially for low-frequency auxin inputs), as quantified by ℳ. The top panel shows 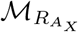, which is less sensitive to changes in *κ* and *P* than 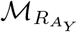 shown in the bottom panel, which characterizes the ARFY transcriptional response. 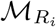

Figure 11 show how certain model parameters change the characteristic response of NAP subnetwork model to different oscillatory auxin stimuli. We deem it possible that similar NAP motifs may act as a filter, operating within a subnetwork to selectively respond to auxin signals of particular frequencies and amplitudes. Such a motif could then activate or deactivate other subnetworks of the NAP in response to auxin signals with particular frequencies.

The NAP controls a cells transcriptional response to particular auxin stimuli. Modelling effort (in this paper, and in the wider literature) has been largely focussed on steady-state behaviour and constant or piecewise-constant auxin stimuli. However, it is clear that certain aspects of auxin signalling closely tied to rhythmic processes in the plant, such as circadian rhythms [46, 9], and so modelling the dynamic response of signalling modules to such non-constant auxin signals is necessary to develop a more complete understanding of the NAP. Here, we have demonstrated a plausible mechanism that governs frequency-selective responses to auxin input using only a few interacting NAP components, and demonstrated that altering only a small subset of parameters results in a wide range of model output—demonstrating the richness of dynamics that can emerge even from relatively simple NAP modules.

## 5 Discussion

Whilst the basic protein-protein and protein-DNA interactions underlying the function of the NAP are well studied, the number of possible model topologies means that a complete model of the entire NAP in a species like *Arabidopsis thaliana* is intractable. Nevertheless, the modelling framework we introduce allows for the exploration of dynamics within specific NAP subnetworks. In this regard, our modelling framework serves as a general-purpose tool that can be used to analyse these dynamics by incorporating knowledge and data concerning specific protein-protein and protein-DNA interactions.

The three examples we provide demonstrate the principle dynamics emerging from three NAP signalling modules. Examples I and II provide examples, built from known and well-studied interactions, that, for certain parameter regimes and ranges of auxin concentrations, show switch-like behaviour in response to changes in auxin concentration. Moreover, we see that if upregulation of the repressor ARF introduced in Example II is (whether by other NAP components or by another pathway altogether), effectively disables switching behaviour.

In contrast, in Example III, the coupled activator ARFs, ARFX and ARFY operate in tandem, but with characteristically different transcriptional responses. Whereas ARFX rapidly responds to increased auxin, ARFY shows a delayed, gradual response (especially when ARFY dimerisation occurs slowly). These two ARFs showed characteristically different transcriptionally responses to oscillatory inputs, too, with ARFX having primarily steady expression levels with small amounts of oscillation, and ARFY having relatively large oscillations. This difference is particularly noticeable for low values of *κ* and auxin oscillations with period greater than 10 hours. In this way, it is possible that target genes have distinct, simultaneous responses to the same auxin stimuli, even though they regulated by the same NAP subnetwork. Such richness may help explain the diversity of temporal response profiles shown in Section 2. Further modelling promises to shed light on the function and robustness of NAP subnetworks containing similar and seemingly redundant ARFs with overlapping targets [16].

Generally, we may expect that certain cells are capable of exhibiting switch-like or graded responses to similar auxin signals depending on a particular extracellular and intracellular context. This is certainly true for the three examples presented above, where bistability exists only in a specific window of auxin concentration and, as demonstrated for Example II, only for specific parameter regimes. These distinct functional modes are surely necessary given the diversity of auxin responses we observe in plant tissue: processes like phototropism [39] and gravitropism [2] direct growth via smooth, graded responses to auxin gradients found in the tissue; whereas bistable, switch-like behaviour drives cell-fate decision-making in processes such as lateral-root formation [22], for example.

### Model limitations

With the aim of producing a parsimonious model for general usage, certain details have been omitted from our treatment of the auxin signalling pathway. Firstly, the model describes only a single form of auxin (most likely IAA), whereas numerous synthetic and endogenous forms exist and have different characteristics. Proper consideration of other auxins would require additional model parameters. However, in the absence of comprehensive data detailing auxin-mediated decay rates for each Aux/IAA, this approach would introduce many more unknown model parameters.

Similarly, binding to the promoter regions is assumed identical across all genes (that is, for each of the mRNA state variables). This is certainly an inaccurate assumption because certain promoter motifs that ARFs bind to are known, and their frequency and location vary greatly across the different ARF and Aux/IAA promoter regions [28].

It would be possible to include such differences of protein-promoter binding kinetics by including a separate Markov model component for each gene. However, this would again introduce a large number of unknown model parameters because there is, currently, little data regarding the association and dissociation rates for different ARFs, Aux/IAAs and dimers thereof.

Furthermore, the chosen ODE-modelling paradigm is limited in that it does not account for stochasticity. The quasi steady-state assumption is valid when promoter-protein binding occurs rapidly compared to other parts of the model, however, if this does not hold, stochasticity could have a significant effect on the output of the model—for example, when there are small numbers of a given signalling component present in the nucleus [37].

Whilst these assumptions and limitations may mask the complexity of some aspects of the underlying biology, our approach allows us to explore possible dynamics of NAP subnetworks, and especially to how these dynamics depend on dimerisation rates. Additional details may be added to the model in future work to address specific questions regarding other aspects of the model, or to integrate insights from experimental data (such as target-gene specific promoter binding dynamics).

### Future work

The analysis presented in Section 2 was collected using a microarray. Such microarrays usually only capture a subset of transcripts from the entire target genome. However, newer next-generation RNA sequencing (RNA-seq) methods allow the entire transcriptome to be analysed in an unbiased manner. To our knowledge, there are no published RNA-seq datasets that can rival the temporal resolution of auxin-modulated gene expression profiles in the same plant tissue provided by this microarray study. The collection and analysis of RNA-seq data from similar experiments with different plant species and tissue types may provide a more complete picture of auxin-signalling dynamics, and allow us to observe behaviour of NAP subnetworks with varying numbers of ARFs and Aux/IAAs.

It is natural to expand this work to consider other NAP subnetworks. In particular, the examples presented above consider only subnetworks with single Aux/IAAs even though it is possible to describe the action of multiple distinct Aux/IAA species in the same framework. A key piece of experimental evidence regarding the potential role of multiple Aux/IAAs is that different members of the Aux/IAA family confer different auxin binding rates [7]. In the examples presented above, we saw how a wide range of responses emerge from systems with only two ARFs (with different preferences for dimerisation) and a single Aux/IAA. We might expect that systems with multiple Aux/IAAs with different binding and decay rates exhibit similar hysteresis and multistability under certain parameter regimes. Further studies should explore similar subnetworks involving distinct Aux/IAA species with different binding and decay rates.

Non-canonical auxin-signalling components [32, 18] (namely, non-canonical ARFs and non-canonical Aux/IAAs) can also be modelled using our approach. For example, in the case of a non-canonical ARF lacking a DNA-binding domain, we can disallow homodimers of this non-canonical ARF from binding to the promoter. Similarly, a non-canonical Aux/IAA which lacks Domain II and so lacks the canonical mechanism for auxin-mediated decay can be modelled simply by setting the relevant auxin-mediated decay rate to zero. NAP subnetworks containing such non-canonical components are attractive for modelling as the additional model complexity is reduced by these missing interactions (in comparison to the addition of a canonical signalling ARF or Aux/IAA).

As mentioned above, the modelling framework introduced here is limited to ARF and Aux/IAA dimers, excluding higher order oligomers from the description. Yet, there is evidence that such complexes can play a functional role *in planta* [38]. Extending our framework would present a significant challenge. In particular, the mere combinatorics of how many complexes can form is prohibitive and would require an in depth investigation, identifying suitable simplifying assumptions to capture the main features whilst being computationally tractable. As we have illustrated above, even dimers alone confer the NAP with a very rich potential for diverse responses to auxin; this additional layer of complexity would surely uncover further interesting behaviour.

Furthermore, other plausible mechanisms of ARF-mediated transcriptional regulation of ARFs (besides dimerisation/multimerisation and direct binding to the promoter) could be explored. For example, it is known that ARF5 indirectly promotes tasi-ARFs which cause degradation of ARF4 [25, 51], affecting the dynamics of the NAP. Hence, we may see indirect repression of ARFs by other promoter ARFs (such as ARF5) through this pathway. Future models should consider such mechanisms of interaction, and the interaction of the NAP with other signalling pathways.

## Supporting information

Article and supplementary material

## Data Accessibility

All codes used for model simulation, analysis, and the production of figures are available on GitHub [40]. The previously-published data analysed in Section 2 is available online at the EBML-EMI data repository [50].

## Acknowledgements

This research was supported by a grant from The Leverhulme Trust. Grant number: RPG-2024-061.

