## Supplementary material for "A general mathematical framework for modelling subnetworks of the nuclear auxin pathway": Article and supplementary material

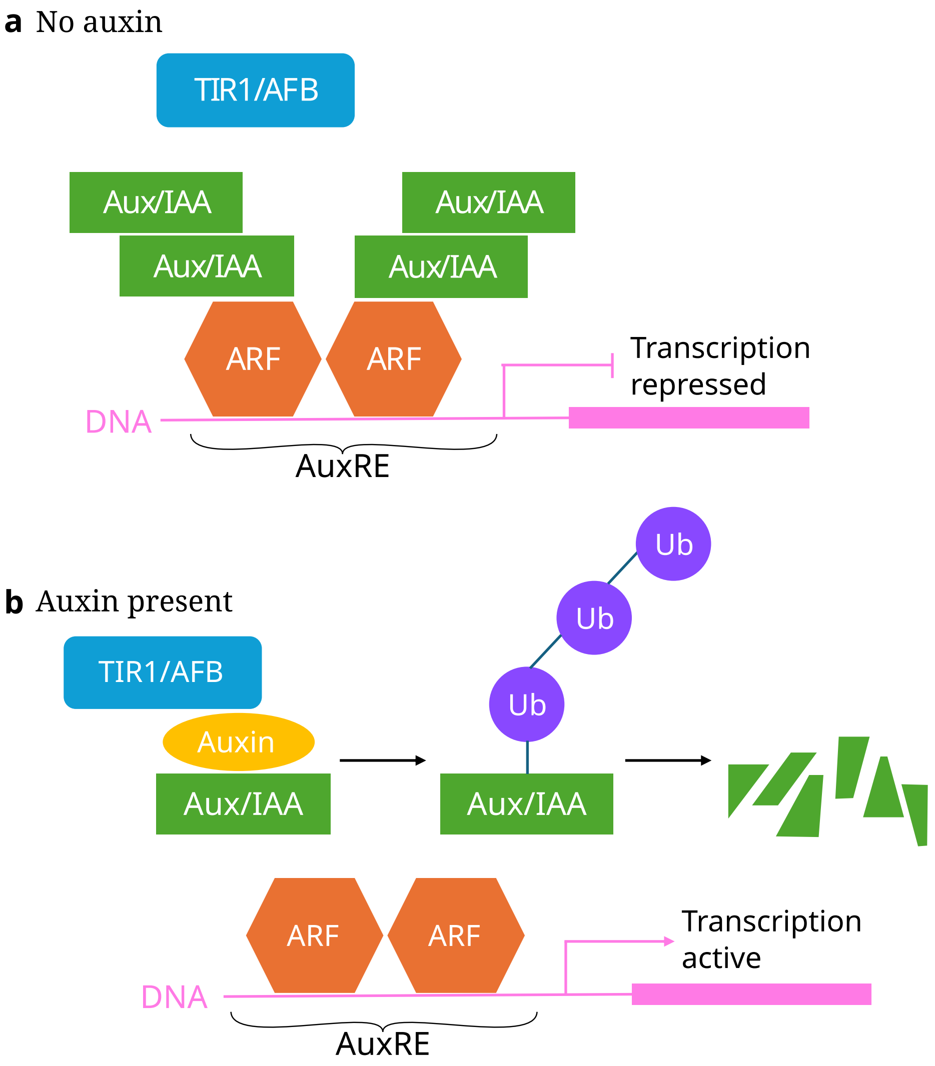


Figure 1: The canonical mechanism by which the NAP causes a transcriptional response in target genes. **a** When there is little or no auxin, Aux/IAAs are free to bind to ARFs (which themselves bind to the DNA) and repress transcription of the target gene. **b**: Auxin causes the ubiquitination and subsequent decay of Aux/IAAs, freeing promoter ARFs to up-regulate the target gene.

To better understand the emergent behaviour arising from such interactions, we propose a mathematical modelling framework that has the flexibility to include any of these possible regulatory

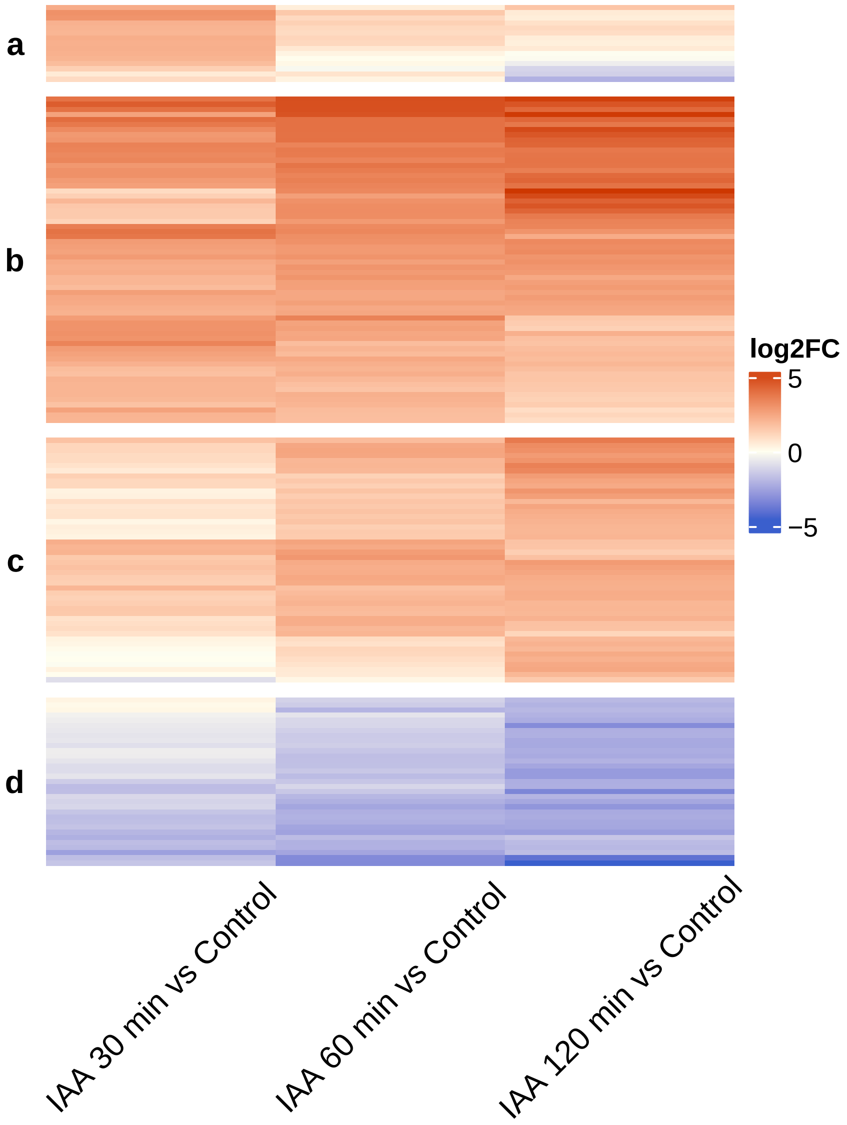


Figure 2: Heatmaps of temporal profiles of gene expression in response to IAA treatment in *Arabidopsis thaliana* root meristems [46]. Genes with a relative differential expression greater than 4 (that is, a $\log_2$-fold change greater than 2) are clustered according to their response profile using Euclidean distance and complete linkage. Each row shows the $\log_2$-fold change (**log2FC**) for a given gene at different time points of IAA treatment relative to the control (no IAA). A positive $\log_2$-fold change (red shades in colour bar) denotes upregulation of a given gene for a given duration of IAA treatment, whereas negative $\log_2$-fold change (blue shades in colour bar) denotes downregulation of a given gene. Genes clustered by $\log_2$-fold change across timepoints roughly fall into four groups showing up-/down-regulation on different timescales. Group **a**: Genes which are rapidly upregulated at 30 minutes, but for which gene expression falls at later time points. Group **b**: Genes which are consistently upregulated across all time points. Group **c**: Genes which are slowly upregulated—often showing little upregulation at 30 minutes, but which become more upregulated at later time points. Group **d**: downregulated genes: some of which are downregulated rather slowly, similar to the delayed upregulation exhibited by group **c**.

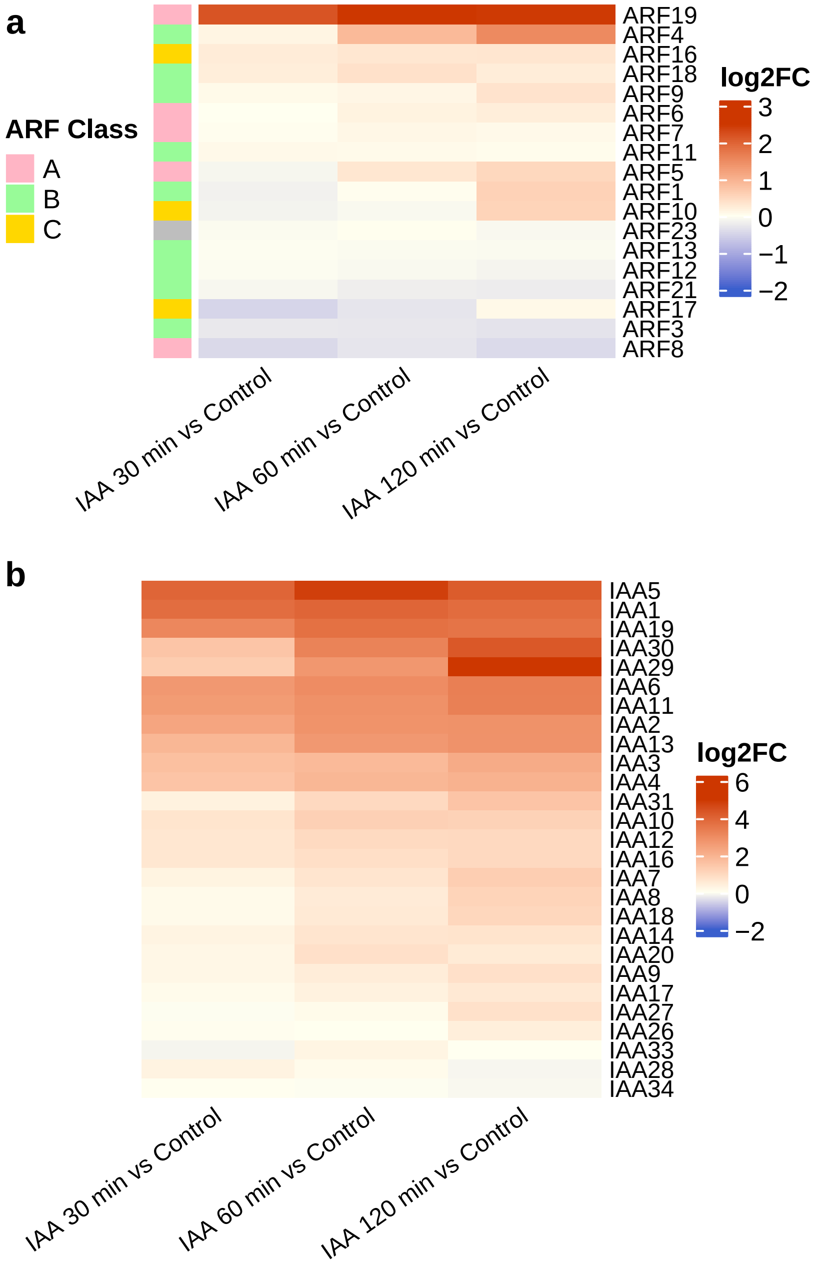


Figure 3: Heatmaps showing relative changes in expression level of *Arabidopsis thaliana* auxin-signalling components after auxin (IAA) treatment. (**a**): $\log_2$ fold-changes in expression of ARFs after IAA treatment compared to a control (no IAA). Genes are clustered according to their response profiles using Euclidean distance and complete linkage. The colours on the left of the heatmap show the class (that is, A, B, or C) of ARF that a given transcript belongs too (ARF23 is coloured grey as it is denoted a pseudogene [8]). Note how a range of different expression patterns can be seen within each ARF class. **b**: $\log_2$ fold-changes in Aux/IAA expression after IAA treatment compared to a control (no IAA). Whilst some Aux/IAAs (IAA28 and IAA34) are slightly down-regulated, the $\log_2$-fold change is much greater than $-2$ (a typical cutoff value).

Moreover, we assume that all reactions between species follow the mass-action law. Then for a single ARF and a single Aux/IAA, we have,

$$\frac{\mathrm{d}A}{\mathrm{d}t} = k_A R_A - d_A A - 2A^2 k_{A,A} - AI k_{A,I} + 2d_{A,A} D_{A,A} + d_{A,I} D_{A,I}\,, \tag{1}$$

$$\frac{\mathrm{d}I}{\mathrm{d}t} = k_I R_I - d_I I - AI k_{A,I} - 2I^2 k_{I,I} + 2d_{I,I} D_{I,I} + d_{A,I} D_{A,I}\,, \tag{2}$$

$$\frac{\mathrm{d}D_{A,A}}{\mathrm{d}t} = k_{A,A} A^2 - d_{A,A} D_{A,A}\,, \tag{3}$$

$$\frac{\mathrm{d}D_{A,I}}{\mathrm{d}t} = k_{A,I} AI - d_{A,I} D_{A,I}\,, \tag{4}$$

$$\frac{\mathrm{d}D_{I,I}}{\mathrm{d}t} = k_{I,I} I^2 - d_{I,I} D_{I,I} \tag{5}$$

where $k_{A,A}$, $k_{A,I}$, and $k_{I,I}$ are association rates; $d_{A,A}$, $d_{A,I}$, and $d_{I,I}$ are the corresponding dissociation rates; and $k_A$ and $d_A$, and $k_I$ and $d_I$ are the production and decay rates for ARFs and Aux/IAA proteins, respectively. The production and decay rates for ARFs, that is $k_A$ and $d_A$, are assumed to be constant.

It is well known that auxin-mediated decay of Aux/IAAs happens via ubiquitination when Aux/IAA is bound to $\mathrm{SCF}^{\mathrm{TIR1}}$ [23]. This degradation pathway is described more completely by previous models [29, 2, 5], whereby the rate of Aux/IAA decay is dependent not only on the concentration of auxin, but also on the concentration of $\mathrm{SCF}^{\mathrm{TIR1}}$. For the sake of simplicity, we omit these details, introducing the variable $a$ to represent the *effective auxin concentration* such that the decay rates for Aux/IAAs are linearly dependent on $a$,

$$d_I = d_0 + d_1 a\,, \tag{6}$$

where $d_0$ is the rate of decay in the absence of auxin, and $d_1$ is a parameter characterising the auxin dependence of Aux/IAA decay.

To capture transcriptional regulation, we let $\mathbf{g}(t)$ be a vector containing the probabilities that the promoter is bound to a particular monomer/dimer at time $t$. Then, we model the binding of ARFs and Aux/IAAs to promoter regions of DNA according to,

$$\frac{\mathrm{d}\mathbf{g}}{\mathrm{d}t} = \frac{1}{\tau_G} \mathbf{Q}_{\mathbf{g}}(\mathbf{x}, \boldsymbol{\theta})^\top \mathbf{g}\,, \tag{7}$$

where $\mathbf{Q}_{\mathbf{g}}$ is a transition-rate matrix which depends on the models state vector $\mathbf{x}$ (containing all state variables) and parameter vector ($\boldsymbol{\theta}$), and $\tau_G$ is some representative timescale that dictates the overall rate of the dynamics, but does not affect the steady state, $\lim_{t\to\infty}\mathbf{x}(t)$. For fixed $\mathbf{x}$ and $\boldsymbol{\theta}$, this portion of the model is a conservative Markov model [41].

$$\frac{\mathrm{d}R_i}{\mathrm{d}t} = K_i(\mathbf{g}; \boldsymbol{\theta}) - d_R R_i \,, \tag{8}$$

where $d_R$ is the assumed constant decay rate of the mRNA (assumed equal for mRNA concentrations corresponding to any ARF or Aux/IAA species); and $K_i(\mathbf{g}, \boldsymbol{\theta})$ is a function which maps the promoter states and parameter vectors to some transcription rate.

Equation (7) describes how the probability that the promoter is in each possible state changes over time. In particular, we denote the components of $\mathbf{g}$ to be $g_{A,A}, g_{A,I}$ and $g_0$, representing the probability that promoter is: bound to an ARF-ARF dimer; bound to an ARF-Aux/IAA dimer; and unbound, respectively. The effective transition rate is then,

$$K_i(\mathbf{x}, \boldsymbol{\theta}) = K_{G_{A,A},i} g_{A,A} + K_{G,i} g_0 + K_{G_{A,I},i} g_{A,I} \,, \tag{9}$$

where $K_{G_{A,A},i}$ is a constant parameter which quantifies the rate of protein translation when an ARF-ARF homodimer is bound to the promoter, $K_{G_{A,I}}$ is the transcription rate of gene $i$ when an ARF-Aux/IAA dimer is bound to the promoter, and $K_{G,i}$ is the transcription rate of gene $i$ due solely to the basal transcription machinery, when the no ARF-ARF or ARF-Aux/IAA dimers are bound to the promoter due solely to the basal transcription machinery (that is, the basal transcription rate). These variables are model parameters and, as such, are components of the model's parameter vector, $\boldsymbol{\theta}$ This scenario is shown by the schematic in Figure 4.

We assume that the promoter-protein dynamics occur rapidly in relation to other parts of the model. Such an assumption is made implicitly in other models, which use probabilistic arguments to define transcription-rate functions [29]. This quasi steady-state assumption allows for the model to be simplified, setting $g_{D_{A,A}}$, $g_{D_{A,I}}$, $g_A$ and $g_0$ by computing the steady-state of Equation (7).

The above-transcription parameters allow a range of activational and repressive effects to be modelled, for every gene represented in the model. For instance, if we consider a single ARF and Aux/IAA, we can omit the subscripts and set $K_{G_{A,A},i} = 1$ and $K_{G,i} = K_{G_{A,I}} = 0$ such that transcription of the $i$ gene only occurs when an ARF-ARF dimer is bound to the promoter. Consequently, the maximal transcription rate is 1, and as the ARF-ARF dimer concentration, $D_{A,A}$, increases, the transcription rate saturates. An example of the relationship between this homodimer concentration, $D_{A,A}$, and a given transcription rate is shown in the Supplementary Material (Section S1). Note that when considering multiple ARFs and Aux/IAAs, these formulas will include additional promoter states and state-dependent transcription rates, and so, the relationship between these concentrations and the effective transcription rates may be more complicated.

Another configuration which shows the flexibility of this formulation is where ARF and Aux/IAA transcription is promoted by ARF-ARF homodimers, but repressed by ARF-IAA heterodimers.

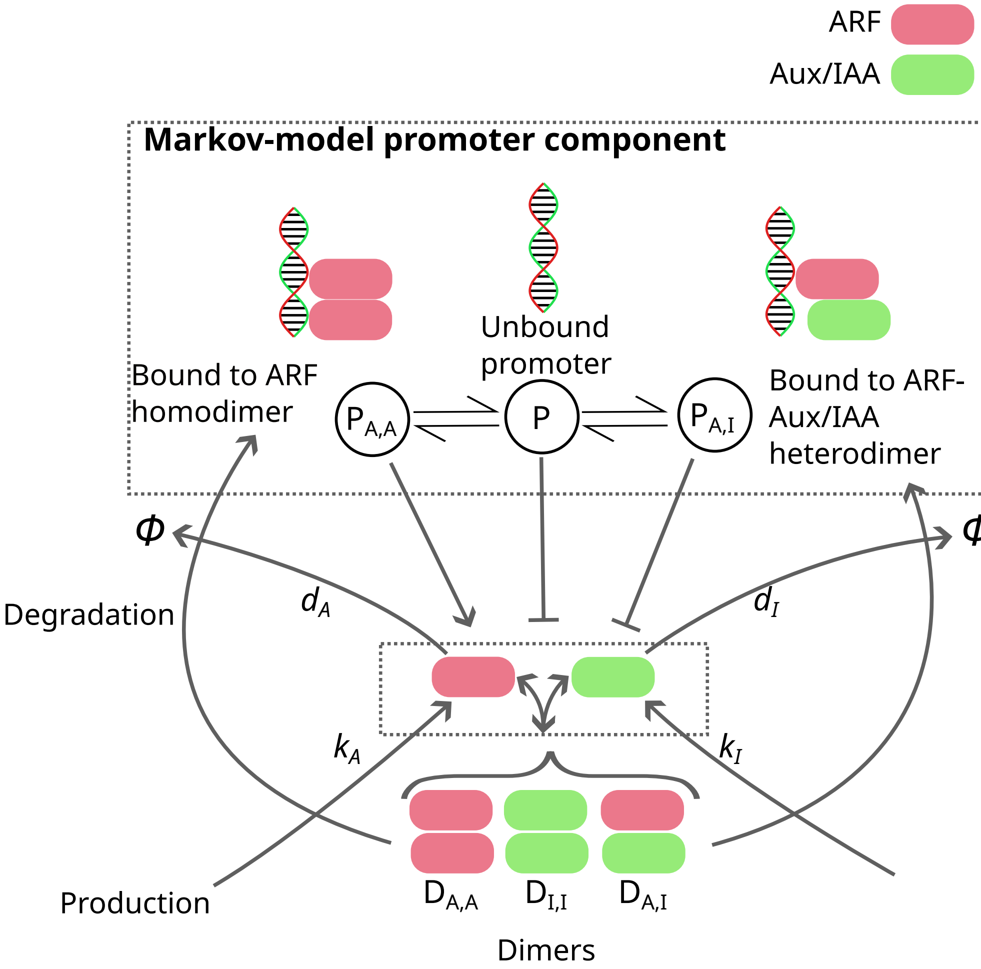


Figure 4: Schematic showing the various components of our NAP modelling framework. A simple, single-ARF, single Aux/IAA NAP is shown for illustrative purposes. The topmost dashed rectangle shows the Markov model describing protein-promoter binding. The probability of the promoter being in each state dictates transcription rates for both ARFs and Aux/IAAs (independently). The lower dashed rectangle shows the production, decay, and dimerisation of ARFs and Aux/IAAs. The concentrations of monomers and dimers affect the transition rates in the Markov model, and hence (indirectly) the transcription rates. Though only a single ARF and Aux/IAA are shown, the same principles can be extended to models with many ARFs and Aux/IAAs. Note that Aux/IAAs lack a DNA-binding domain like that of an ARF, and as such are unable to bind directly to the DNA.

This is achieved by ensuring a positive basal transition rate, $K_{G,i} > 0$ for each gene $i$ (that is, basal transcription of ARF and Aux/IAA). Then we choose $K_{G_{A,A},i} > K_{G,i}$ such that ARF-ARF homodimers promote the transcription of both genes, and $K_{G_{A,I},i} < K_{G,i}$, such that ARF-Aux/IAA dimers repress transcription.

In any case, the transcription rate is dependent on the dimer concentrations $D_{A,A}$ and $D_{A,I}$, which are governed by Equations (3) and (4). As $\tau_G \to 0$, the timescale of protein-promoter dynamics decreases. For small $\tau_G$ we can approximate the system using a *quasi steady-state assumption*,

$$\frac{\mathrm{d}\mathbf{y}}{\mathrm{d}t} = \mathbf{f}(\mathbf{y}, t; \boldsymbol{\theta}), \tag{10}$$

$$\mathbf{g}(t) = \mathbf{g}_\infty(\mathbf{y}, \boldsymbol{\theta}), \tag{11}$$

where $\mathbf{f}(y, t; p)$ is the vector-valued function describing the evolution of the state variables as in Equation (1)–(5), and the function $\mathbf{g}_\infty$ maps these state variables, denotes the equilibrium point of the promoter Markov model corresponding to the chosen parameter values, $\boldsymbol{\theta}$, and the instantaneous state of the model $\mathbf{y}$ (the concentrations of protein complexes in particular). This equilibrium distribution is determined by finding the non-trivial solution to Equation (7) with elements summing to one, that is, $\mathbf{1}^\top \mathbf{g} = 1$. If the protein-promoter dynamics are assumed to happen quickly (compared to other dynamics in the model), the resulting ODE system computed explicitly is "stiff", meaning that significant computational work is required to simulate the model. This problem is alleviated by the quasi steady-state formulation. Reducing the number of state variables in this way also simplifies the numerical analyses presented in the following sections [41].

#### 3.2 General modelling framework for multiple ARFs and Aux/IAAs

We now provide notation for NAP models containing multiple ARFs. Examples II and III below are models of this form, containing two distinct ARFs and a single Aux/IAA species. Because it is simple to extend this modelling framework to an arbitrary number of ARFs and Aux/IAAs, we provide the notation and software tools required to define and analyse such models. However, we do not consider any modles with multiple Aux/IAA in this article. Nevertheless, we introduce the necessary notation.

We use $A_1, A_2, \ldots, A_N$ to denote the concentration of $N$ different ARF species. Similarly, we notate concentrations of the various Aux/IAA species as $I, I_2, \ldots, I_M$ where $M$ is the number of Aux/IAA species represented in the model. Then, we denote a dimer of the $i^{\text{th}}$ ARF transcript with the $j^{\text{th}}$ ARF transcript as $D_{A_i,A_j}$. Likewise, $D_{A_i,I_j}$ denotes the concentration of dimers formed from ARF $i$ and Aux/IAA $j$, and $D_{I_i,I_j}$ denotes the concentration of dimers formed from Aux/IAA $i$ and Aux/IAA $j$. As before, we assume that the order of the indices of the dimer are inconsequential.

Using this notation we can describe the dynamics of NAP with multiple ARF and/or Aux/IAA species under the modelling assumptions listed above. For example, for a model including two distinct ARF species, but only a single Aux/IAA species, we have,

$$
\begin{aligned}
\frac{\mathrm{d}A_1}{\mathrm{d}t} =& k_A R_{A_1} - d_{A_1} A_1 - 2{A_1}^2 k_{A_1,A_1} + 2 d_{A_1,A_1} D_{A_1,A_1} - k_{A_1,I} A_1 I + d_{A_1,I} D_{A_1,I} \\
& - k_{A_1,A_2} A_1 A_2 - d_{A_1,A_2} D_{A_1,A_2} \,, && (12) \\
\frac{\mathrm{d}A_2}{\mathrm{d}t} =& k_A R_{A_2} - d_{A_2} A_2 - 2{A_2}^2 k_{A_2,A_2} + 2 d_{A_2,A_2} D_{A_2,A_2} - k_{A_2,I} A_2 I + d_{A_2 I} D_{A_2,I} \\
& - k_{A_1,A_2} A_1 A_2 - d_{A_1,A_2} D_{A_1,A_2} \,, && (13) \\
\frac{\mathrm{d}I}{\mathrm{d}t} =& k_I R_I - d_I I - A_1 I k_{A_1,I} - A_2 I k_{A_2,I} - 2 I^2 k_{I,I} + 2 d_{I,I} D_{I,I} + d_{A_1,I} D_{A_1,I} \\
& + d_{A_2,I} D_{A_2,I} \,, && (14) \\
\frac{\mathrm{d}D_{A_1,A_1}}{\mathrm{d}t} =& {A_1}^2 k_{A_1,A_1} - d_{A_1,A_1} D_{A_1,A_1} \,, && (15) \\
\frac{\mathrm{d}D_{A_1,A_2}}{\mathrm{d}t} =& A_1 A_2 k_{A_1,A_2} - d_{A_1,A_2} D_{A_1,A_2} \,, && (16) \\
\frac{\mathrm{d}D_{A_2,A_2}}{\mathrm{d}t} =& {A_2}^2 k_{A_2,A_2} - d_{A_2,A_2} D_{A_2,A_2} \,, && (17) \\
\frac{\mathrm{d}D_{A_1,I}}{\mathrm{d}t} =& A I k_{A_1,I} - d_{A_1,I} D_{A_1,I} \,, && (18) \\
\frac{\mathrm{d}D_{A_2,I}}{\mathrm{d}t} =& A_2 I k_{A_1,I} - d_{A_2,I} D_{A_2,I} \,, && (19) \\
\frac{\mathrm{d}D_{I,I}}{\mathrm{d}t} =& I^2 k_{I,I} - d_{I,I} D_{I,I} && (20)
\end{aligned}
$$

As in the single-ARF, single-Aux/IAA model, the concentration of each species of mRNA is governed by Equation (8), and the corresponding effective transcription rates computing via the same Markov-model formulation as in Equations (7) and (9). Moreover, we again simplify this model by omitting the monomer-bound promoter states and using a quasi steady-state assumption (as in Equations (10)–(11)), where protein-promoter binding events are assumed to occur much more rapidly than other processes in the model. The resulting quasi steady-state of the promoter Markov model then determines transcription rates similarly to Equation (9), but with additional terms for the additional promoter states.

For the examples provided in this section, the model is further simplified by the assumption that lone ARFs can not be bound directly to the promoter, and that dimerisation of a promoter-bound monomer is not possible. In this case, the Markov chain described by Equation (7) has a simple "hub-and-spoke" topology (where each promoter-bound state is directly connected only to the unbound state), such that the steady-state probabilities are given by,

$$
\begin{aligned}
g_{A_i,A_j} =& \frac{\phi_{A_i,A_j}}{1 + \sum_{k,l} \phi_{A_k,A_l} + \sum_{k,l} \phi_{A_k,I_l}} \,, && (21) \\
g_{A_i,I_j} =& \frac{\phi_{A_i,I_j}}{1 + \sum_{k,l} \phi_{A_k,A_l} + \sum_{k,l} \phi_{A_k,I_l}} \,, && (22) \\
\text{and } g_0 =& \frac{1}{1 + \sum_{k,l} \phi_{A_k,A_l} + \sum_{k,l} \phi_{A_k,I_l}} \,, && (23)
\end{aligned}
$$

where $g_{A_j,A_k} g_{A_i,I_j}, g_{A_j}$ and $g_0$ denote the probability that the promoter is in each of the given states, and $\phi_{A_i,A_j}$ and $\phi_{A_i,I_j}$ denote the ratio of association and dissociation rates for ARF-dimer-bound promoter states, and ARF-Aux/IAA-dimer-bound states, respectively. Note how this quasi steady-state assumption effectively reduces the number of necessary parameters, as the above steady-states may be expressed simply in terms of ratios association and dissociation rates.

Accordingly, for a model with $N$ ARF species and $M$ Aux/IAA species, we may write the effective transcription rate by combining the state-dependent transcription rates for each of the promoter states (with occupation probabilities denoted, as above, by $g_{A_i,A_j}, g_{A_i,I_j}$ and $g_0$),

$$K_i(\mathbf{x},\boldsymbol{\theta}) = \big(\sum_{j=1}^{N}\sum_{k=j}^{N} g_{A_j,A_k} K_{G_{A_j,A_k},i}\big) + \big(\sum_{j=1}^{N}\sum_{k=1}^{M} g_{A_j,I_k} K_{G_{A_j,A_k},i}\big) + g_0 K_{G,i}\,. \tag{24}$$

Note that this formulation could be extended as to also include monomer-bound promoter states. However, this is not necessary for the work presented here.

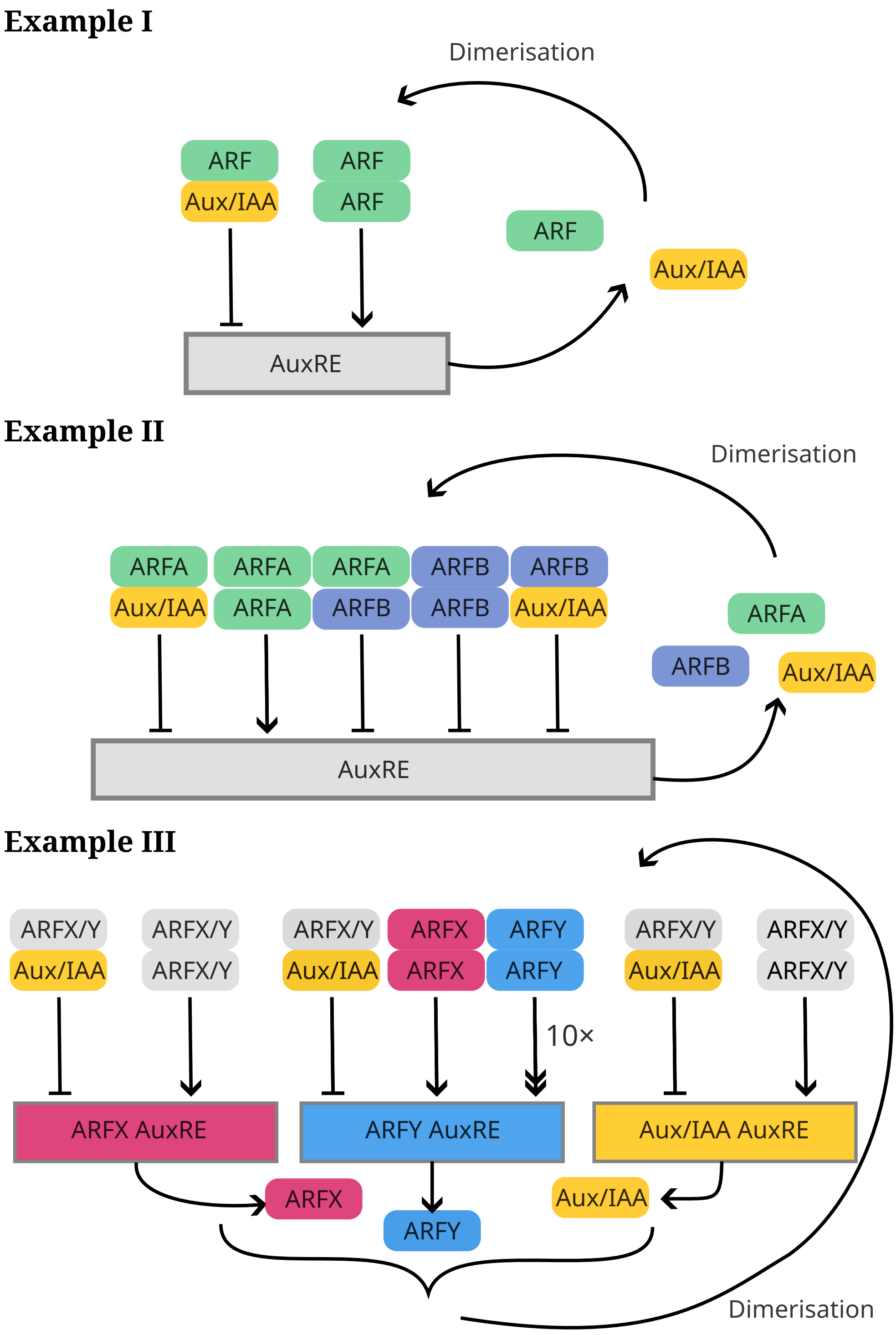


Figure 5: Diagrams of the example NAP subnetworks explored in this article. **Example I**: A single-ARF, single-Aux/IAA model where transcription (of ARF and Aux/IAA genes) is promoted by ARF homodimers, but repressed by ARF-Aux/IAA heterodimers. **Example II**: the previous model but with an additional repressor ARF with mRNA transcribed at a constant rate unless the promoter is bound to a complex containing Aux/IAA (not pictured). **Example III**: A double-ARF feed-forward motif in which two class-A ARFs (ARFX and ARFY) promote each other and where one ARF, ARFY, promotes itself particularly strongly.

To capture the promotion and co-repression dynamics described above, we set the state-dependent transcription rates to be,

$$K_{G_{A,A},R_A} = 1.0\,, \tag{25}$$
$$K_{G_{A,I},R_A} = 0.0\,, \tag{26}$$
$$K_{G,R_A} = 0.001\,, \tag{27}$$

and similarly, those for BDL to be,

$$K_{G_{A,A},R_I} = 1.0\,, \tag{28}$$
$$K_{G_{A,I},R_I} = 0.0\,, \tag{29}$$
$$K_{G,R_I} = 0.015\,, \tag{30}$$
$$\tag{31}$$

where the first subscript (a dimer) corresponds to the complex bound to the promoter region, and the second subscript denotes the type of mRNA being produced—MP ($A$) or BDL ($I$); the basal transcription rates where no compound is bound to the promoter are denoted by $K_{G,A}$ and $K_{G,I}$ for MP and BDL, respectively.

From these state-dependent transcription rates, we see that an increase in BDL-MP dimers represses transcription of MP and BDL because this (rapidly) increases the probability that an MP-BDL dimer is bound to the DNA, and $K_{G,A} > K_{G_{A,I},A}$ and $K_{G,I} > K_{G_{A,I},I}$. Likewise, we see that an increase in ARF-ARF dimers effects an increase in transcription of the same genes because $K_{G,A} < K_{G_{A,A},A}$ and $K_{G,I} < K_{G_{A,A},I}$.

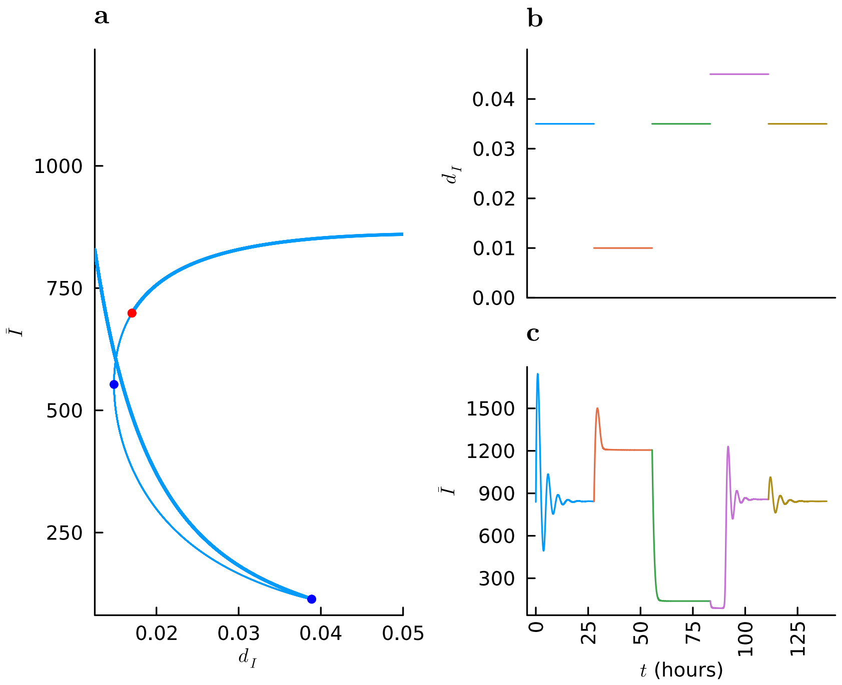


Figure 6: Multistability in the NAP can lead to hysteretic responses for certain auxin concentrations, as shown by the total (bound and unbound) Aux/IAA concentration, $\bar{I} = I + 2D_{I,I} + D_{A,I}$. Three different auxin concentrations are applied to the model, and we see that a drop in auxin sends the model to a sole stable branch with high $\bar{I}$ (shown in Panel **a**), and the application high auxin concentration returns the network to the original branch. For auxin intermediate auxin concentrations, however, the system may converge to one of two stable equilibria. Panel **a**: bifurcation diagram showing possible steady states; stable steady states are marked by the thicker blue line and unstable steady states by a thinner line; blue circles show the location of bifurcation points and the red circle denotes a Hopf bifurcation. Panel **b**: a piecewise constant auxin input signal. Panel **c**: the model's response to the above auxin signal in $\bar{I}$.

$$K_{G_{A_A,A_A},R_{A_B}} = \lambda, \tag{32}$$
$$K_{G_{A_A,A_B},R_{A_B}} = \lambda, \tag{33}$$
$$K_{G_{A_A,I},R_{A_B}} = 0, \tag{34}$$
$$K_{G_{A_B,A_B},R_{A_B}} = \lambda, \tag{35}$$
$$K_{G_{A_B,I},R_{A_B}} = 0, \tag{36}$$
$$K_{G,R_{A_B}} = \lambda \tag{37}$$

where $\lambda$ is the aforementioned transcription rate.

The repressive effect of the newly-added ARF is achieved by assuming no transcription of ARFA ($R_{A_A}$) or BDL ($R_I$) whenever the promoter is bound to a dimer containing ARFB. That is,

$$K_{G_{A_A,A_B},R_{A_A}} = 0, \tag{38}$$
$$K_{G_{A_B,A_B},R_{A_A}} = 0, \tag{39}$$
$$K_{G_{A_B,I},R_{A_A}} = 0, \tag{40}$$
$$K_{G_{A_A,A_B},R_I} = 0, \tag{41}$$
$$K_{G_{A_B,A_B},R_I} = 0, \tag{42}$$
$$K_{G_{A_B,I},R_I} = 0, \tag{43}$$
$$K_{G,R_{A_B}} = 0. \tag{44}$$

In this way, an increased ARFB concentration (say from $\lambda$ being increased) leads to reduced transcription of MP ($R_{A_A}$) and BDL ($R_I$), as expected. Increasing $A_B$ also leads to increased formation of $D_{A_A,A_B}$ dimers, reducing $A_A$ and hence $D_{A_A,A_A}$. Again, a full table of model parameter values is provided in the Supplementary Material.

In Figure 7, we see that this system has a bifurcation point and bistability provided the ARFB transcription rate (in the case that no dimer containing Aux/IAA is bound to the promoter), $\lambda$, is not too large. In other words, we see that an increase in repressor-ARF activity can eliminate bistability.

The above example shows how action of a repressor ARF might be described mathematically, and shows a possible mechanism for the deactivation of any given signalling motif in the NAP. A repressor ARF (ARFB in this example) could be transcriptionally regulated in many ways (by other parts of the NAP, or by other signalling networks entirely) modifying the transcription rate, $\lambda$. In this way, certain subnetworks (operating as logic units such as binary switches), might be be activated or deactivated according to environmental and developmental cues.

Example I showed that a single-ARF, single-Aux/IAA network could act as either a bistable or monostable circuit depending on parameters. By including a repressor ARF, we observe that there are a greater number of scenarios in which the network output can switch between bistable and monostable regimes. This switch can happen independently of parameters present in Example I (including auxin concentration) and can be obtained solely by altering regulation of the repressor ARF. This capability would mean that external cues regulating a repressor ARF could modulate the outcome of a subnetwork containing this component to provide a response distinct from other subnetworks without that ARF.

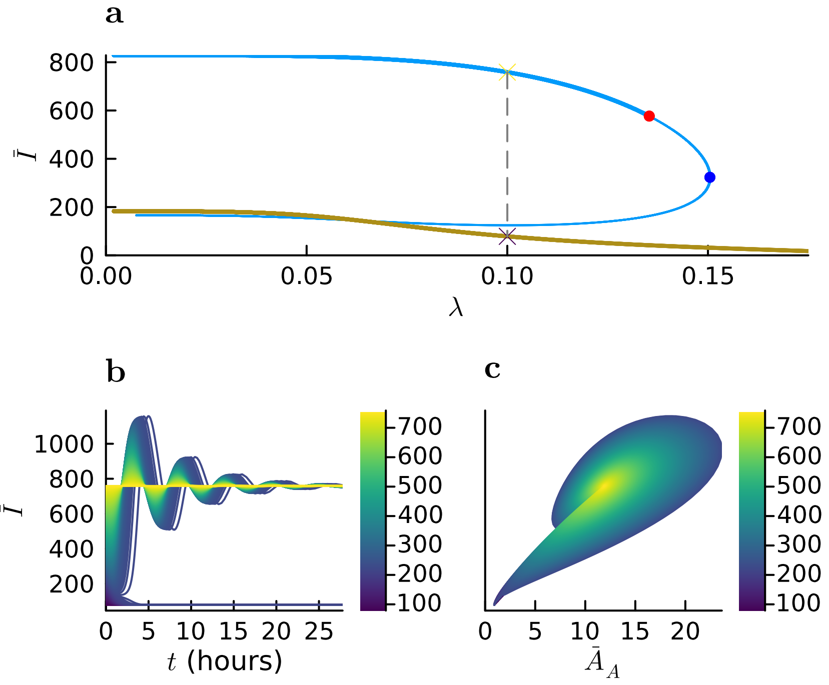


Figure 7: Provided repressor-ARF activity is not excessive, the MP-BDL model (augmented with a repressor ARF) can be multistable. The total BDL concentration (both bound and unbound), $\bar{I}$ is shown on the y-axis. **a** shows a bifurcation diagram with a bifurcation point and bistability between two stable branches (shown in thick blue and brown lines) for various values of the transcription-rate $\lambda$. The thin blue line shows a branch of unstable equilibria. A subcritical Hopf bifurcation is displayed in red, and a branch point is shown in blue. **b**: simulations under the same parameter set using various initial conditions selected along the dashed line segment shown in Panel **a**. Panel **c**: phase-plane plot of the same simulated trajectories, showing total BDL concentration (bound and unbound), $\bar{I}$, vs. total MP concentration, $\bar{A}$.

Then, for ARFY we suppose that the transcription only occurs when the promoter is bound to an ARF-ARF dimer, that is, $K_{G_{A_X,A_X},R_{A_Y}} = 0.2$, $K_{G_{A_X,A_Y},R_{A_Y}} = 0.2$, $K_{G_{A_Y,A_Y},R_{A_Y}} = 2.5$, and all other promoter-dependent ARFY transition rates (including the basal transcription rate) are zero: $K_{G_{A_i,I},R_{A_Y}} = K_{G,R_{A_Y}} = 0$ for $i \in \{1, 2\}$. In this way, ARFY self-promotes and is promoted (less strongly) by ARFX, and repressed by Aux/IAA. As for previous sections, complete tables of model parameters and initial conditions may be found in the Supplementary Material.

A maximal ARFY transcription rate is obtained when the probability of an ARFY-ARFY dimer being bound to the promoter is 1, that is, when there is a large amount of ARFY-ARFY dimers present (both in absolute terms, and relatively when compared to the other compounds). However, we also expect there to be low amounts of ARFY present in a low-auxin environment, because the basal transcription rate is 0. Therefore, we may expect a response to high-auxin concentration starting with a small rapid increase as ARFX-ARFX dimers bind to the promoter (increasing ARFX and ARFY transcription). Following this, depending on the speed of ARFY dimerisation, we may see a slower sustained increase as more ARFY is produced and more ARFY-ARFY dimers form and bind to the promoter region. A full table of model parameters is provided in the Supplementary Material.

Under these assumptions, certain functional characteristics of this network may be modulated by modifying association/dissociation rates of particular dimers. In particular, we fix the ratio of association and dissociation rates for ARFY dimerisation with itself, other ARFs, and Aux/IAA, scaling these rates together. That is, we introduce a scaling factor $\kappa$, and update the association/dissociation rates for ARFY-ARFY dimerisation as follows,

$$\begin{aligned} k'_{A_X,A_Y} &= \frac{1}{\kappa} k_{A_X,A_Y}\,, \\ d'_{A_X,A_Y} &= \frac{1}{\kappa} d_{A_X,A_Y}\,. \end{aligned} \tag{45}$$

$$\begin{aligned} k'_{A_Y,A_Y} &= \frac{1}{\kappa} k_{A_Y,A_Y}\,, \\ d'_{A_Y,A_Y} &= \frac{1}{\kappa} d_{A_Y,A_Y}\,. \end{aligned} \tag{46}$$

$$\begin{aligned} k'_{A_Y,I} &= \frac{1}{\kappa} k_{A_Y,I}\,, \\ d'_{A_Y,I} &= \frac{1}{\kappa} d_{A_Y,I}\,. \end{aligned} \tag{47}$$

(48)

In this way, increasing $\kappa$ slows down the dimerisation of ARFY with other compounds (including itself), without affecting the model's equilibrium concentrations (as demonstrated in the Supplementary Material B). Only the transient behaviour of the model is affected, causing changes in the time-taken for ARFX and ARFY to respond to auxin stimulus, as demonstrated in the following section.

Broadly speaking, as the effective auxin concentration, $a$, increases, the total Aux/IAA concentration, $\bar{I} = I + D_{A_X,I} + D_{A_Y,I} + 2D_{I,I}$ (including both bound and unbound forms) decreases, though for some auxin concentrations the model exhibits multi-stability, as shown by the bifurcation diagram in Figures 8 (panels **a** and **c**). Aside from affecting steady-states, parameters have a large impact on the transient response of the model to different auxin stimuli. We proceed to explore how dimer association and dissociation rates affect the time taken to produce transcriptional responses to step-changes in auxin concentration. Then, we consider how these same parameters affect characteristics of the models' response to oscillatory auxin stimuli.

##### 4.3.1 Auxin response time

Figure 8 shows how larger values of $\kappa$ introduces a delay to the auxin-induced upregulation of ARFY under this model. Whilst the transcriptional response of ARFX varies little under these changes in $\kappa$, the time taken for the upregulation of ARFY is controlled by this parameter.

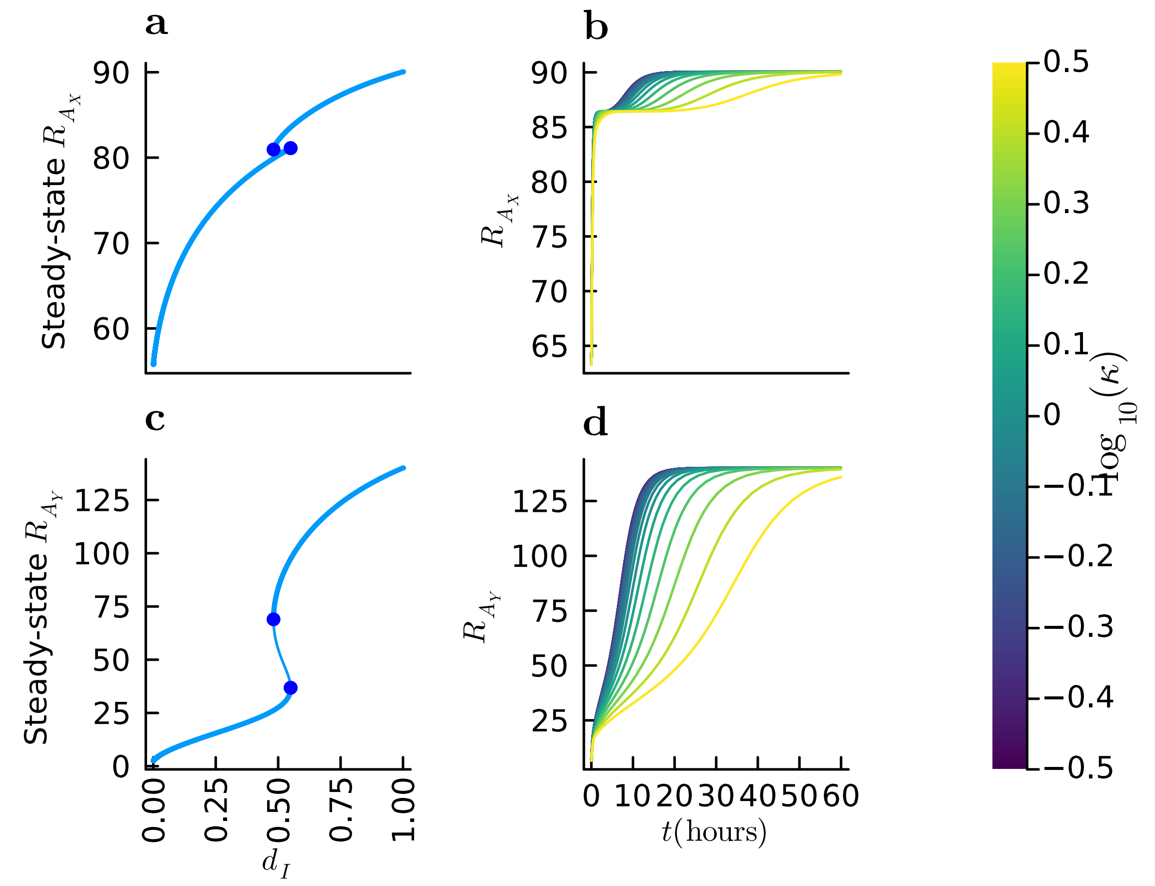


Figure 8: Transcriptional response to auxin treatment in $R_{A_X}$ and $R_{A_Y}$ shown for various values of $\kappa$ found by equilibrating the model under low-auxin conditions and simulating under high auxin. **a**: bifurcation diagram showing steady states, denoted by blue circles, as the auxin concencentration, $a$, varies. **b**: a selection of ARFX-mRNA trajectories obtained by simulating the model with different values of $\kappa$. Panels **c** and **d**: show similar plots for ARFY mRNA, that is $R_{A_Y}$. For high $\kappa$, ARFY dimerisation occurs slowly, causing a delayed response. Whereas, the speed of transcriptional response for ARFX varies little under different values of $\kappa$.

##### 4.3.2 Response to sinusoidal auxin concentration

Temporally oscillating auxin signals and fluctuations are found in plants with various periods—the root clock [30], for example, or auxin rhythmic centrifugal waves in the shoot [15]. It is also known that there is interplay between the nuclear NAP and cell-level auxin transport processes (such as transport via PIN and AUX/LAX proteins). Accordingly, we expand our attention beyond constant auxin concentrations and consider the application of a sinusoidal auxin stimulus. We model a representative oscillatory signal by fixing our model's intracellular auxin concentration to a sinusoidal function, $a(t) = \frac{1}{2} + \frac{1}{2}\sin(\frac{t}{2P\pi})$, where $P$ is the period of the oscillation. An example simulation of the model is shown in Figure 9.

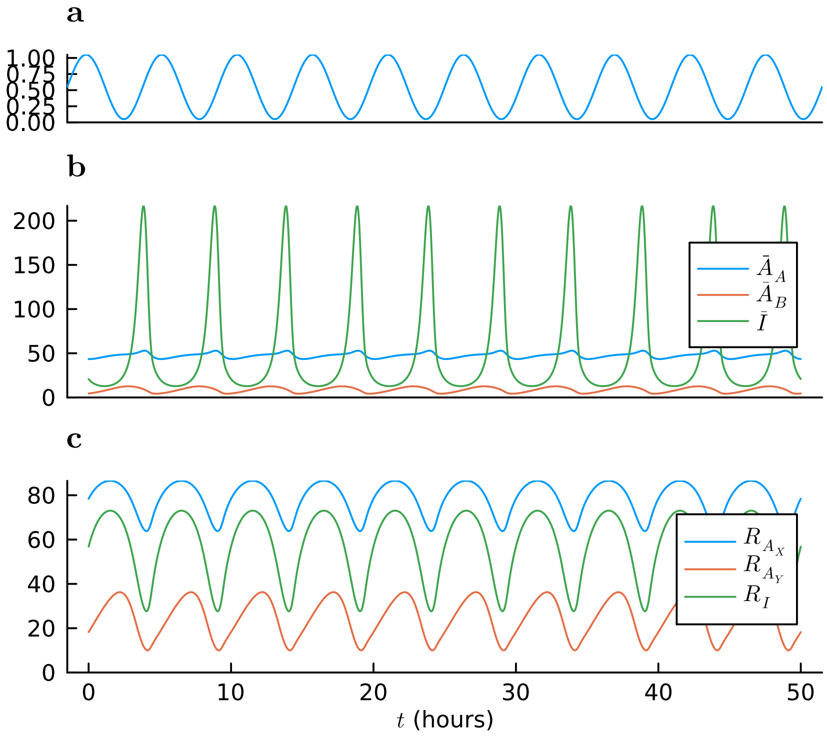


Figure 9: Output of the simulated model when $\kappa = 100$ and $P = 5$hours. The Aux/IAA concentration rapidly increases and then decreases during the portion of the signal where auxin is low. This, through its binding with ARFX and ARFY, prompts a transcriptional response for ARFX, ARFY and Aux/IAA. However, these transcriptional responses have different characteristics: $R_{A_X}$ has a roughly steady-state expression when compared to $R_{A_Y}$ which exhibits relatively larger oscillations. Panel **a** the auxin signal with period $P = 5$ hours. Panel **b**: total protein concentration of all complexes containing ARFA, ARFB and Aux/IAA ($\bar{A}_A, \bar{A}_B, \bar{I}$, respectively). Panel **c**: mRNA concentrations for the genes coding for ARFX, ARFY and Aux/IAA ($R_{A_X}, R_{A_Y}$, and $R_I$, respectively).

To characterise the resulting mRNA output we compute,

$$\mathcal{M}_{R_i} = \frac{1}{10P \max_{t\in[t_0,t_0+10P]}(R_i(t))} \int_{t_0}^{t_0+10P} R_i \, \mathrm{d}t \,, \tag{49}$$

where $P$ is the period of the auxin input and $R$ is some mRNA concentration, and where $R_i$ is the time-course of interest, being an mRNA concentration such as $R_{A_X}, R_{A_Y}$, or $R_I$. In words, $\mathcal{M}_{R_i}$ indicates whether the resulting signal is oscillatory or not. For a constant signal, we would have $\mathcal{M}_{R_i} = 1$, and for a signal consisting of short, rapid spikes we would have $\mathcal{M}_{R_i} \approx 1$. Since we wish to characterise the long-term behaviour of the model at its limit cycle, so we compute $\mathcal{M}_{R_i}$ by simulating the model until $t = 1,000P$, and computing $\mathcal{M}_{R_i}$ using $t_0 = 990P$.

Figure 11 demonstrates how the functional output of the model depends on $\kappa$ and the period of the input signal (auxin concentration). Certain frequencies of auxin input lead to qualitatively different ARFY transcriptional responses provided a suitable value of $\kappa$. This demonstrates how the dimerisation dynamics of ARFs in the model may give rise to different temporal expression profiles for target genes in response to non-constant auxin signals.

A heatmap showing $\mathcal{M}_{R_i}$ for both ARFX and ARFY, and various $(\kappa, P)$ for both ARFX and ARFY is shown in Figure 11. Little difference is found in $\mathcal{M}_{R_i}$ for ARFY across our selection of $(\kappa, P)$ pairs. However, there are noticeable differences in the behaviour of $R_{A_Y}$ as shown in more detail in Figures 10, where $R_{A_Y}$ is plotted under numerous simulations using various $(\kappa, P)$ values. Here, we see that the maximal ARFY response in each oscillation is greater for smaller values of $\kappa$ and that the position of these maxima is shifted when compared to the high-$\kappa$ trajectories.

We summarise the mRNA concentrations for both ARFs in Figure 11. In this example, the rate of ARFY dimerisation controls the sensitivity of the model output to $P$, the period of the auxin input. The output signal is most sensitivity (as quantified by Equation (49)) to changes in the auxin-signal period when dimerisation is rapid (that is, for small $\kappa$). Notably, the expression profiles of ARFX are much more uniform than those of ARFY across different values of $\kappa$.

Figure 11 show how certain model parameters change the characteristic response of NAP subnetwork model to different oscillatory auxin stimuli. We deem it possible that similar NAP motifs may act as a filter, operating within a subnetwork to selectively respond to auxin signals of particular frequencies and amplitudes. Such a motif could then activate or deactivate other subnetworks of the NAP in response to auxin signals with particular frequencies.

The NAP controls a cells transcriptional response to particular auxin stimuli. Modelling effort (in this paper, and in the wider literature) has been largely focussed on steady-state behaviour and constant or piecewise-constant auxin stimuli. However, it is clear that certain aspects of auxin signalling closely tied to rhythmic processes in the plant, such as circadian rhythms [46, 9], and so modelling the dynamic response of signalling modules to such non-constant auxin signals is necessary to develop a more complete understanding of the NAP. Here, we have demonstrated a plausible mechanism that governs frequency-selective responses to auxin input using only a few interacting NAP components, and demonstrated that altering only a small subset of parameters results in a wide range of model output—demonstrating the richness of dynamics that can emerge even from relatively simple NAP modules.

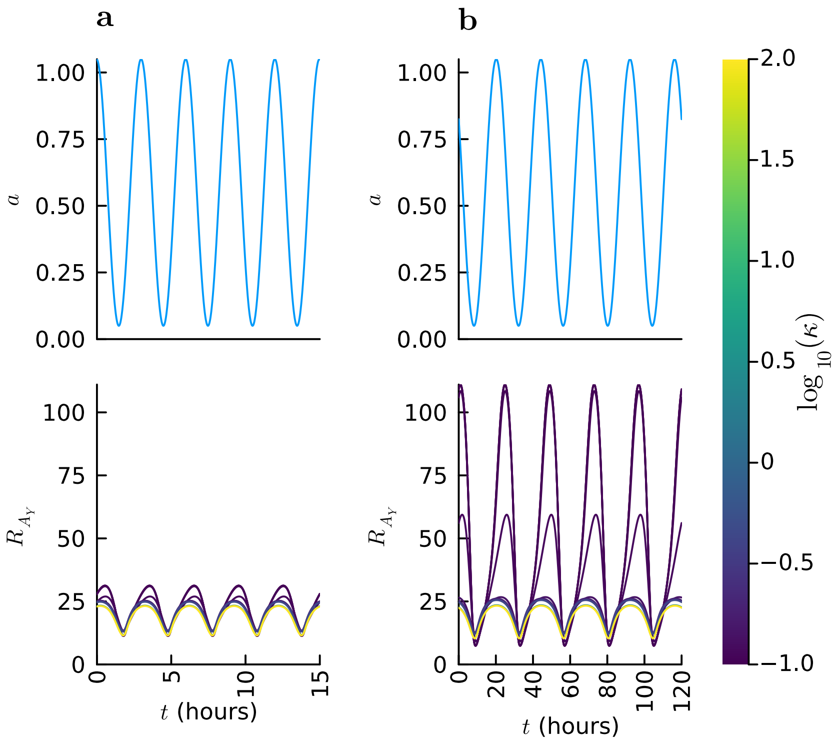


Figure 10: Normalised trajectories for two different auxin-input periods. Column **a**: simulation of a signal with period $P = 3$ hours. Column **b**: simulation of a signal with period $P = 24$ hours. Note that not enough periods have been simulated for the model to reach steady state in some cases. Here we see that the relative size of oscillations depends on $\kappa$, with larger $\kappa$ leading to relatively small oscillations, and smaller $\kappa$ leading to relatively large oscillations. This effect is particularly noticeable for lower-frequency stimuli (that is, for larger $P$).

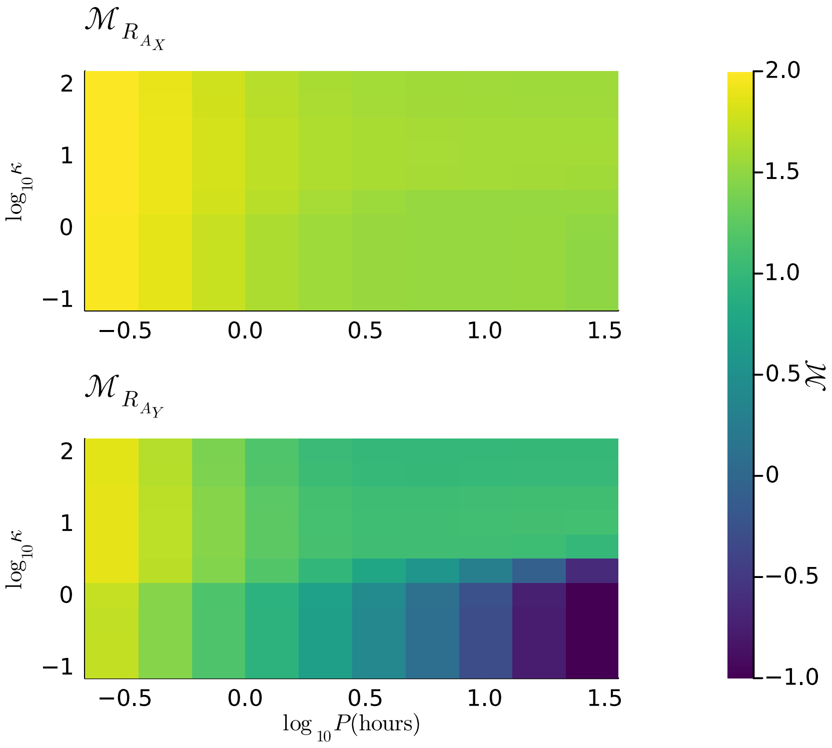


Figure 11: A heatmap showing $\mathcal{M}_{R_i}$ for various combinations of $\kappa$ and $P$ for both ARFX mRNA and ARFY mRNA, with $\kappa \approx 1$ signifying constant expression levels. The ARFY transcriptional response to auxin signals with different inputs is largely dependent on the timescale of ARFY dimerisation (especially for low-frequency auxin inputs), as quantified by $\mathcal{M}$. The top panel shows $\mathcal{M}_{R_{A_X}}$, which is less sensitive to changes in $\kappa$ and $P$ than $\mathcal{M}_{R_{A_Y}}$ shown in the bottom panel, which characterises the ARFY transcriptional response. $\mathcal{M}_{R_i}$

**Acknowledgements** This research was supported by a grant from The Leverhulme Trust. Grant number: RPG-2024-061.

### S1 State-dependent transcription model

Figure S1 shows the relationship between $D_{A,A}$ and the effective transcription rate for two genes in a simple, single-ARF, single Aux/IAA model, under a representative parameter set. Here, we set all dimer-protein association and dissociation rates to 1.0 and set the state-dependent transcription rates as follows: $K_{G_{A,A},R_A} = K_{G,R_A} = 1.0$, $K_{G_{A,I},R_A} = K_{G_{A,I},R_I} = 0.0$, $K_{G_{A,A},R_A} = 2$, and $K_{G,R_I} = \frac{1}{2}$. The only state variables relevant in Figure S1 are the two ARF-containing dimer concentrations: $D_{A,I}$ (set to 1) and $D_{A,A}$ (the abscissa).

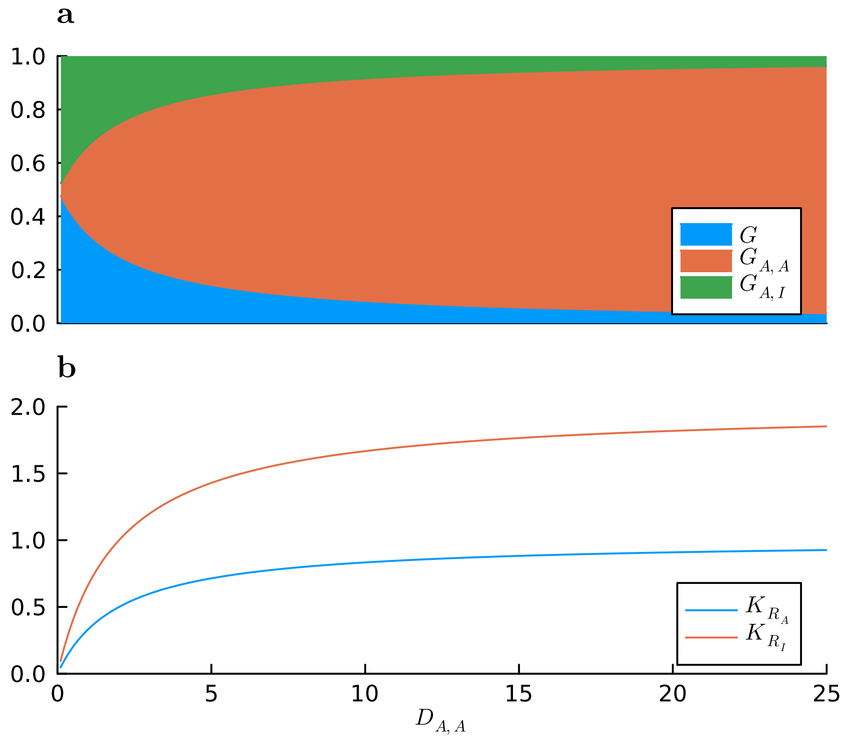


Figure S1: For a promoter ARF, $A$, we expect that transcription increases as the dimer concentration, $D_{A,A}$ increases. **a**: the probability that the promoter is in each state (bound to an ARF-ARF dimer, bound to an ARF-Aux/IAA dimer, or unbound) at time $t$. **b**: the resultant transcription rate for the two genes, both of which are transcribed only when an ARF-Aux/IAA dimer is not bound to the promoter (that is, in the ARF homodimer and unbound states).

### S2 Model parameters

A complete table of model parameters used in Example I (Section 44.1) are shown in Table S1. Note that the bifurcation parameter explored in Figure 6 is the BDL decay rate, $d_I$, and since BDL decays rapidly in the presence of auxin, we associate high auxin concentrations with greater $d_I$, and low auxin concentrations with small $d_I$.

| Parameter | Value | Description |
|---|---|---|
| **Protein production** | | |
| $k_A$ | 1.0 | Rate of ARF production from RNA |
| $k_I$ | 1.0 | Rate of IAA production from RNA |
| **Protein decay** | | |
| $d_I$ | 0.1 | Decay rate of Aux/IAAs |
| $d_{I_x}$ | 10.0 | Rate of auxin-mediated decay of Aux/IAA |
| $d_A$ | 1.0 | Decay rate of ARF proteins |
| **mRNA decay** | | |
| $d_R$ | 1e-4 | RNA decay rate |
| **Dimerisation** | | |
| $d_{D_{I,I}}$ | 1.0 | Dissociation rate of dimer $D_{I,I}$ |
| $k_{D_{A,I}}$ | 5.0 | Association rate of dimer $D_{A,I}$ |
| $d_{D_{A,I}}$ | 1.0 | Dissociation rate of dimer $D_{A,I}$ |
| $k_{D_{I,I}}$ | 5.0 | Association rate of dimer $D_{I,I}$ |
| $d_{D_{A,A}}$ | 1.0 | Dissociation rate of dimer $D_{A,A}$ |
| $k_{D_{A,A}}$ | 5.0 | Association rate of dimer $D_{A,A}$ |
| **Transcription** | | |
| $K_{G_{A,I},R_I}$ | 0.0 | Rate of transcription for $R_I$ for promoter $G_{A,I}$ |
| $K_{G_{R_I}}$ | 1.5e-4 | Rate of transcription for $R_I$ for promoter $G$ |
| $K_{G_{A,I},R_A}$ | 0.0 | Rate of transcription for $R_A$ for promoter $G_{A,I}$ |
| $K_{G,R_A}$ | 7.5e-6 | Rate of transcription for $R_A$ for promoter $G$ |
| $K_{G_{A,A},R_A}$ | 0.01 | Rate of transcription for $R_A$ for promoter $G_{A,A}$ |
| $K_{G_{A,A},R_I}$ | 0.01 | Rate of transcription for $R_I$ for promoter $G_{A,A}$ |

Table S1: Full model parameters for Example I

Full model parameters for Example II are provided in Table S2. Note that these parameters are largely the same as those in Table S1, and that the ARFB transcription-rate parameter, $\lambda$, is not included here as it is not given a fixed value. Similarly, a complete table of parameters for Example III is provided in Table S3.

| Variable | Value | Description |
|---|---|---|
| **Protein production** | | |
| $k_A$ | 1.0 | Rate of ARF production from RNA |
| $k_I$ | 1.0 | Rate of IAA production from RNA |
| **mRNA decay** | | |
| $d_R$ | 0.001 | RNA decay rate |
| **Protein decay** | | |
| $d_I$ | 0.03 | Decay rate of Aux/IAAs |
| $d_A$ | 1.0 | Decay rate of ARF proteins |
| **Dimerisation** | | |
| $d_{D_{A_2,I}}$ | 1.0 | Dissociation rate of dimer $D_{A_2,I}$ |
| $d_{D_{A_1,A_2}}$ | 0.05 | Dissociation rate of dimer $D_{A_1,A_2}$ |
| $k_{D_{A_1,A_1}}$ | 5.0 | Association rate of dimer $D_{A_1,A_1}$ |
| $d_{D_{I,I}}$ | 1.0 | Dissociation rate of dimer $D_{I,I}$ |
| $k_{D_{A_2,A_2}}$ | 5.0 | Association rate of dimer $D_{A_2,A_2}$ |
| $k_{D_{A_2,I}}$ | 5.0 | Association rate of dimer $D_{A_2,I}$ |
| $k_{D_{A_1,I}}$ | 5.0 | Association rate of dimer $D_{A_1,I}$ |
| $d_{D_{A_1,I}}$ | 1.0 | Dissociation rate of dimer $D_{A_1,I}$ |
| $k_{D_{I,I}}$ | 5.0 | Association rate of dimer $D_{I,I}$ |
| $d_{D_{A_2,A_2}}$ | 1.0 | Dissociation rate of dimer $D_{A_2,A_2}$ |
| $d_{D_{A_1,A_1}}$ | 1.0 | Dissociation rate of dimer $D_{A_1,A_1}$ |
| $k_{D_{A_1,A_2}}$ | 0.05 | Association rate of dimer $D_{A_1,A_2}$ |
| **Transcription** | | |
| $K_{G_{A_2,A_2},R_{A_1}}$ | 0.0 | Rate of transcription for $R_{A_1}$ for promoter $G_{A_2,A_2}$ |
| $K_{G_{A_2,I}R_{A_2}}$ | 0.0 | Rate of transcription for $R_{A_2}$ for promoter $G_{A_2,I}$ |
| $K_{G_{A_1,I},R_{A_1}}$ | 0.0 | Rate of transcription for $R_{A_1}$ for promoter $G_{A_1,I}$ |
| $K_{G_{A_2,I},R_{A_1}}$ | 0.0 | Rate of transcription for $R_{A_1}$ for promoter $G_{A_2,I}$ |
| $K_{G_{A_2,A_2},R_{A_2}}$ | 1.0 | Rate of transcription for $R_{A_2}$ for promoter $G_{A_2,A_2}$ |
| $K_{G_{A_1,A_1},R_{A_1}}$ | 0.01 | Rate of transcription for $R_{A_1}$ for promoter $G_{A_1,A_1}$ |
| $K_{G_{A_1,A_1},R_{A_2}}$ | 1.0 | Rate of transcription for $R_{A_2}$ for promoter $G_{A_1,A_1}$ |
| $K_{G_{A_1,I},R_{A_2}}$ | 0.0 | Rate of transcription for $R_{A_2}$ for promoter $G_{A_1,I}$ |
| $K_{G_{A_1,A_1},R_I}$ | 0.01 | Rate of transcription for $R_I$ for promoter $G_{A_1,A_1}$ |
| $K_{G_{A_1,A_2}R_{A_2}}$ | 1.0 | Rate of transcription for $R_{A_2}$ for promoter $G_{A_1,A_2}$ |
| $K_{G_{A_1,A_2},R_I}$ | 0.0 | Rate of transcription for $R_I$ for promoter $G_{A_1,A_2}$ |
| $K_{G,R_{A_1}}$ | 7.5e-6 | Rate of transcription for $R_{A_1}$ for promoter $G$ |
| $K_{G_{A_2,I},R_I}$ | 0.0 | Rate of transcription for $R_I$ for promoter $G_{A_2,I}$ |
| $K_{G,R_{A_2}}$ | 1.0 | Rate of transcription for $R_{A_2}$ for promoter $G$ |
| $K_{G,R_I}$ | 1.5e-4 | Rate of transcription for $R_I$ for promoter $G$ |
| $K_{G_{A_2,A_2},R_I}$ | 0.0 | Rate of transcription for $R_I$ for promoter $G_{A_2,A_2}$ |
| $K_{G_{A_1,A_2},R_{A_1}}$ | 0.0 | Rate of transcription for $R_{A_1}$ for promoter $G_{A_1,A_2}$ |
| $K_{G_{A_1,I},R_I}$ | 0.0 | Rate of transcription for $R_I$ for promoter $G_{A_1,I}$ |

Table S2: Full model parameters for Example II

| Variable | Value | Description |
|---|---|---|
| **Production and decay rates** | | |
| $k_A$ | 0.005 | Rate of ARF production from RNA |
| $k_I$ | 0.1 | Rate of IAA production from RNA |
| $d_R$ | 0.01 | RNA decay rate |
| $d_I$ | 0.05 | decay rate of Aux/IAAs |
| $d_{all}$ | 0.1 | decay rate of ARF proteins |
| **Dimerisation** | | |
| $d_{D_{A_2,I_1}}$ | 0.1 | Dissociation rate of dimer $D_{A_2,I_1}$ |
| $d_{D_{A_1,A_2}}$ | 0.1 | Dissociation rate of dimer $D_{A_1,A_2}$ |
| $k_{D_{A_1,A_1}}$ | 0.05 | Association rate of dimer $D_{A_1,A_1}$ |
| $d_{D_{I_1,I_1}}$ | 0.1 | Dissociation rate of dimer $D_{I_1,I_1}$ |
| $k_{D_{A_2,A_2}}$ | 0.1 | Association rate of dimer $D_{A_2,A_2}$ |
| $k_{D_{A_2,I_1}}$ | 0.1 | Association rate of dimer $D_{A_2,I_1}$ |
| $k_{D_{A_1,I_1}}$ | 0.1 | Association rate of dimer $D_{A_1,I_1}$ |
| $d_{D_{A_1,I_1}}$ | 0.1 | Dissociation rate of dimer $D_{A_1,I_1}$ |
| $k_{D_{I_1,I_1}}$ | 0.1 | Association rate of dimer $D_{I_1,I_1}$ |
| $d_{D_{A_2,A_2}}$ | 0.1 | Dissociation rate of dimer $D_{A_2,A_2}$ |
| $d_{D_{A_1,A_1}}$ | 0.05 | Dissociation rate of dimer $D_{A_1,A_1}$ |
| $k_{D_{A_1,A_2}}$ | 0.01 | Association rate of dimer $D_{A_1,A_2}$ |
| **Transcription** | | |
| $K_{G_{A_2,A_2,R,A_1}}$ | 1.0 | Rate of transcription for $R_{A_1}$ for promoter $G_{A_2,A_2}$ |
| $K_{G_{A_1,I_1,R,A_1}}$ | 0.5 | Rate of transcription for $R_{A_1}$ for promoter $G_{A_1,I_1}$ |
| $K_{G_{A_2,I_1,R,A_1}}$ | 0.5 | Rate of transcription for $R_{A_1}$ for promoter $G_{A_2,I_1}$ |
| $K_{G_{A_1,A_1,R,A_1}}$ | 1.0 | Rate of transcription for $R_{A_1}$ for promoter $G_{A_1,A_1}$ |
| $K_{G_{R,A_1}}$ | 1.0 | Rate of transcription for $R_{A_1}$ for promoter $G$ |
| $K_{G_{A_1,A_2,R,A_1}}$ | 1.0 | Rate of transcription for $R_{A_1}$ for promoter $G_{A_1,A_2}$ |
| $K_{G_{A_2,I_1,R,A_2}}$ | 0.0 | Rate of transcription for $R_{A_2}$ for promoter $G_{A_2,I_1}$ |
| $K_{G_{A_2,A_2,R,A_2}}$ | 2.5 | Rate of transcription for $R_{A_2}$ for promoter $G_{A_2,A_2}$ |
| $K_{G_{A_1,A_1,R,A_2}}$ | 0.25 | Rate of transcription for $R_{A_2}$ for promoter $G_{A_1,A_1}$ |
| $K_{G_{A_1,I_1,R,A_2}}$ | 0.0 | Rate of transcription for $R_{A_2}$ for promoter $G_{A_1,I_1}$ |
| $K_{G_{A_1,A_1},R_{I_1}}$ | 1.0 | Rate of transcription for $R_{I_1}$ for promoter $G_{A_1,A_1}$ |
| $K_{G_{A_1,A_2},R_{A_2}}$ | 0.25 | Rate of transcription for $R_{A_2}$ for promoter $G_{A_1,A_2}$ |
| $K_{G_{A_1,A_2},R_{I_1}}$ | 1.0 | Rate of transcription for $R_{I_1}$ for promoter $G_{A_1,A_2}$ |
| $K_{G_{A_2,I_1},R_{I_1}}$ | 0.0 | Rate of transcription for $R_{I_1}$ for promoter $G_{A_2,I_1}$ |
| $K_{G,R_{A_2}}$ | 0.0 | Rate of transcription for $R_{A_2}$ for promoter $G$ |
| $K_{G_{A_2,A_2},R_{I_1}}$ | 1.0 | Rate of transcription for $R_{I_1}$ for promoter $G_{A_2,A_2}$ |
| $K_{G_{A_1,I_1},R_{I_1}}$ | 0.0 | Rate of transcription for $R_{I_1}$ for promoter $G_{A_1,I_1}$ |
| $K_{G,R_{I_1}}$ | 1.0 | Rate of transcription for $R_{I_1}$ for promoter $G$ |

Table S3: Full model parameters for Example III

### S3 Example III: steady states are independent of $\kappa$

In Section 44.3 of the main text, we introduce model describing a simple NAP subnetwork consisting a single Aux/IAA and two ARFs, ARFX and ARFY. We then introduce a hyperparameter $\kappa$ which controls the relative rate of dimerisation for ARFY. Recall from the main text that the concentrations of proteins and dimers in this model are governed by the equations,

$$\begin{aligned}
\frac{\mathrm{d}A_X}{\mathrm{d}t} =& k_{A_X} - d_{A_X}A_X - 2k_{A_X,A_X}{A_X}^2 + 2d_{A_X,A_X}D_{A_X,A_X} - k_{A_X,I}A_XI + d_{A_X,I}D_{A_X,I} \\
&- k_{A_X,A_Y}A_XA_Y - d_{A_X,A_Y}D_{A_X,A_Y}\,, && \text{(S1)}\\
\frac{\mathrm{d}A_Y}{\mathrm{d}t} =& k_{A_Y} - d_{A_Y}A_Y - 2k_{A_Y,A_Y}{A_Y}^2 + 2d_{A_Y,A_Y}D_{A_Y,A_Y} - k_{A_Y,I}A_YI + d_{A_YI}D_{A_Y,I} \\
&- k_{A_X,A_Y}A_XA_Y - d_{A_X,A_Y}D_{A_X,A_Y}\,, && \text{(S2)}\\
\frac{\mathrm{d}I}{\mathrm{d}t} =& k_I - d_II - k_{A_X,I}A_XI - k_{A_Y,I}A_YI - 2I^2k_{I,I} + 2d_{I,I}D_{I,I} + d_{A_X,I}D_{A_X,I} \\
&+ d_{A_Y,I}D_{A_Y,I}\,, && \text{(S3)}\\
\frac{\mathrm{d}D_{A_X,A_X}}{\mathrm{d}t} =& k_{A_X,A_X}{A_X}^2 - d_{A_X,A_X}D_{A_X,A_X}\,, && \text{(S4)}\\
\frac{\mathrm{d}D_{A_X,A_Y}}{\mathrm{d}t} =& k_{A_X,A_Y}A_XA_Y - d_{A_X,A_Y}D_{A_X,A_Y}\,, && \text{(S5)}\\
\frac{\mathrm{d}D_{A_Y,A_Y}}{\mathrm{d}t} =& k_{A_Y,A_Y}{A_Y}^2 - d_{A_Y,A_Y}D_{A_Y,A_Y}\,, && \text{(S6)}\\
\frac{\mathrm{d}D_{A_X,I}}{\mathrm{d}t} =& k_{A_X,I}AI - d_{A_X,I}D_{A_X,I}\,, && \text{(S7)}\\
\frac{\mathrm{d}D_{A_Y,I}}{\mathrm{d}t} =& k_{A_X,I}A_YI - d_{A_Y,I}D_{A_Y,I}\,, && \text{(S8)}\\
\frac{\mathrm{d}D_{I,I}}{\mathrm{d}t} =& k_{I,I}I^2 - d_{I,I}D_{I,I} && \text{(S9)}
\end{aligned}$$

where each $k_{i,j}$ is an association rate, each $d_{i,j}$ is a dimerisation rate, and each $k_{A_i}$ and $k_{I_i}$ is a production rate depending on the corresponding mRNA concentration. If we assume a steady state, then we have $\frac{\mathrm{d}\mathbf{x}}{\mathrm{d}t} = 0$ where $\mathbf{x}$ is a "state-variables vector" containing the above variables and also the mRNA concentrations. Consequently, the production rates ($k_{A_X}$, $k_{A_Y}$ and $k_I$) are constant, too.

Now, consider the total ARFX concentration obtained by the summation of all ARFX dimers in the system as well as the monomer concentration, $A_X$ itself. Denoting this quantity by $\bar{A_X}$, we have,

$$\bar{A_X} = A_X + 2D_{A_X,A_X} + D_{A_X,I} + D_{A_X,A_Y}\,, \tag{S10}$$

where the factor of two appears owing to the fact that each ARFX homodimer contains two ARFX proteins.

Then we have,

$$\begin{aligned}
\frac{\mathrm{d}\bar{A_X}}{\mathrm{d}t} =& \frac{\mathrm{d}A_X}{\mathrm{d}t} + 2\frac{\mathrm{d}D_{A_X,A_X}}{\mathrm{d}t} + \frac{\mathrm{d}D_{A_X,I}}{\mathrm{d}t} + \frac{\mathrm{d}D_{A_X,A_Y}}{\mathrm{d}t} && \text{(S11)}\\
=& k_{A_X} - A_Xd_{A_X}\,. && \text{(S12)}
\end{aligned}$$

and so, at steady-state we necessarily have,

$$\frac{\mathrm{d}\bar{A}}{\mathrm{d}t} = k_{A_X} - A_Xd_{A_X} = 0\,, \tag{S13}$$

and hence $A_X = \frac{k_{A_X}}{d_{A_X}}$.

Now, consider $\frac{\mathrm{d}}{\mathrm{d}t} D_{A_X,A_X}$. Assuming equilibrium,

$$0 = \frac{\mathrm{d}D_{A_X,A_X}}{\mathrm{d}t} \tag{S14}$$
$$= {A_X}^2 k_{A_X,A_X} - d_{A_X,A_X} D_{A_X,A_X} \tag{S15}$$
$$= \frac{k_{A_X}^2 k_{A_X,A_X}}{d_{A_X}^2} - d_{A_X,A_X} D_{A_X,A_X} \tag{S16}$$

and so, we have

$$D_{A_X,A_X} = \frac{k_{A_X}^2 k_{A_X,A_X}}{d_{A_X}^2 d_{A_X,A_X}}. \tag{S17}$$

Similarly, at steady state, we have

$$D_{I,I} = \frac{k_I^2 k_{I,I}}{d_I^2 d_{I,I}}, \tag{S18}$$

and,

$$D_{A_Y,A_Y} = \frac{k_{A_Y}^2 k_{A_Y,A_Y}}{d_{A_Y}^2 d_{A_Y,A_Y}}. \tag{S19}$$

Similar manipulations yield

$$D_{A_X,A_Y} = \frac{k_{A_X} k_{A_Y} k_{A_X,A_Y}}{d_{A_X} d_{A_Y} d_{A_X,A_Y}} \tag{S20}$$
$$D_{A_X,I} = \frac{k_{A_X} k_I k_{A_X,I}}{d_{A_X} d_I d_{A_X,I}} \tag{S21}$$
$$\text{and } \; D_{A_Y,I} = \frac{k_{A_Y} k_I k_{A_Y,I}}{d_{A_Y} d_I d_{A_Y,I}} \tag{S22}$$

Recall that in Example III we scale $k_{A_Y,A_Y}$ and $d_{A_Y,A_Y}$ identically, keeping $\dfrac{k_{A_Y,A_Y}}{d_{A_Y,A_Y}}$ constant. We can see that the above steady-state concentrations are unaffected by this parameter scaling. Nevertheless, that the transient behaviour observed when it is not at equilibrium varies significantly, as demonstrated in Section 44.3.
